# A specific and portable gene expression program underlies antigen archiving by lymphatic endothelial cells

**DOI:** 10.1101/2024.04.01.587647

**Authors:** Ryan M. Sheridan, Thu A. Doan, Cormac Lucas, Tadg S. Forward, Aspen Uecker-Martin, Thomas E. Morrison, Jay R. Hesselberth, Beth A. Jirón Tamburini

## Abstract

Antigens from protein subunit vaccination traffic from the tissue to the draining lymph node, either passively via the lymph or carried by dendritic cells at the local injection site. Lymph node (LN) lymphatic endothelial cells (LEC) actively acquire and archive foreign antigens, and archived antigen can be released during subsequent inflammatory stimulus to improve immune responses. Here, we answer questions about how LECs achieve durable antigen archiving and whether there are transcriptional signatures associated with LECs containing high levels of antigen. We used single cell sequencing in dissociated LN tissue to quantify antigen levels in LEC and dendritic cell populations at multiple timepoints after immunization, and used machine learning to define a unique transcriptional program within archiving LECs that can predict LEC archiving capacity in independent data sets. Finally, we validated this modeling, showing we could predict antigen archiving from a transcriptional dataset of CHIKV infected mice and demonstrated *in vivo* the accuracy of our prediction. Collectively, our findings establish a unique transcriptional program in LECs that promotes antigen archiving that can be translated to other systems.

## Introduction

Viral antigens persist within lymph nodes following viral infection^1–5^. Persisting antigen, in the case of influenza virus infection, can recruit memory T cells, providing protection against future infections with the same virus^1–3, 5–7^, demonstrating that persisting viral antigens may be an important mediator of immune memory responses. We demonstrated that protein subunit immunization, administered with a TLR agonist subcutaneously, also initiates persistence of the protein antigen in the draining lymph node (dLN)^8–11^. Similar to virus-derived persisting antigen, the vaccine-derived persisting antigen improved T cell memory^8, 10^. We termed this process of durable antigen retention “antigen archiving”^10^.

Lymphatic endothelial cells (LEC)s interact with antigens and virus as they traffic through lymphatic capillaries to lymphatic collectors before entering the lymph node sinus and the fibroblastic reticular cell (FRC) conduit system. These two cell types, LEC and FRC, along with blood endothelial cells (BEC), comprise a larger non-hematopoietic subset of cells within the LN called lymph node stromal cells (LNSC). Many new LNSC subtypes have been defined using single-cell RNA-seq^11–14^, propelling our understanding of the diversity of LNSCs and their contributions to lymph node (LN) function. Despite these recent advances, the functional roles of draining LNSC subpopulations are poorly defined. To begin to understand how LNSCs archive antigens, we conjugated a protein antigen to a stabilized DNA tag and tracked its distribution after immunization using single cell sequencing, quantitatively measuring antigen-DNA levels across different LEC subsets^11^.

While LECs can present peripheral tissue antigens to promote peripheral tolerance, archived antigens are exchanged to migratory conventional dendritic cells (cDC1 and 2) to promote immunity. Consistent with live imaging and antigen presentation assays, antigen-DNA conjugates were only found in CCR7hi migratory cDCs^11^. Thus, as migratory DCs traffic from the skin to the draining lymph node, they interact with antigen bearing LECs, acquiring antigens to improve the effector function of memory CD8 T cells^9–11^. Indeed, some antigen is acquired from LECs that undergo apoptosis during LN contraction^8, 9^. To understand whether this process of LEC apoptosis or LEC-DC antigen exchange could be manipulated, we determined the consequences of immune insult would be to antigen held within LECs. We found that a subsequent inflammatory stimulus within the archiving timeframe yielded increased effector responses compared to those that did not have a secondary stimulus. These improved responses were independent of bystander activation but depended on TCR stimulation^8^. Protection from these boosted memory CD8+ T cells was durable and robust, establishing a model wherein cell-mediated immunity can be manipulated by the presentation of previously archived antigens. Collectively, these studies demonstrate a novel role for the lymph node stromal cells (LNSC) and particularly LECs, in promoting protective T cell immunity at late time points after immunization.

LECs acquire multiple types of protein antigens^8–10^ including purified chikungunya viral E2 glycoprotein, albumin, ovalbumin, Sars-Co-V2 receptor binding domain (RBD)^8^, influenza nucleoprotein (NP), herpes simplex virus glycoprotein B (HSVgB), and vaccinia viral proteins post-infection^10^. In addition to acquisition and retention of foreign protein antigens, LECs may support viral replication, such as Kaposi’s sarcoma associated herpes virus^15^. Recently, single cell mRNA sequencing of LNSCs during CHIKV infection indicated that subsets of LECs that express the scavenger receptor MARCO may support CHIKV RNA replication^16^, consistent with another study showing MARCO-dependent internalization of CHIKV by LN LECs^17^. CHIKV RNA is also detectable in FRCs, which express the CHIKV entry receptor Mxra8^18, 19^. Although CHIKV infection causes LN disorganization and impaired CD8^+^ T cell responses^17, 20^, exploiting this mechanism of viral targeting could lead to a better understanding of the functions of distinct LNSCs. Confirmation of these entry mechanisms used single cell RNA sequencing to detect very low levels of CHIKV RNA within macrophage receptor with collagenous structure (MARCO)-expressing LECs^16, 17^.

In our previous studies of antigen-DNA conjugates, antigen levels were assessed 2 days and 2 weeks post-immunization ^11^. More recently^8^, we found that the protective capacity of archived antigens was maintained at longer timepoints (>2 weeks post-immunization), especially when there was a secondary inflammatory stimulus that caused the release of archived antigens to stimulate memory T cells. However, there are still unanswered questions about how LNSCs archive antigens, including the duration and maintenance of archived antigens within LECs, whether LECs acquire and retain multiple antigens following sequential immunizations, and whether LNSC are intrinsically programmed to archive antigens, or whether they learn to archive additional antigens after acquisition.

## Results

### Antigen persists in discrete cell populations within the lymph node

To study the dissemination of antigens in the draining lymph node after vaccination, we previously developed a single-cell RNA-seq (scRNA-seq) approach to quantify antigen levels *in vivo*^10^ involving conjugation of a model antigen, ovalbumin (ova), to a DNA barcode containing phosphorothioate linkages (psDNA). Using this tool, we showed that immunization with ova-psDNA and vaccinia virus (VV) induces robust antigen archiving^9, 11^. We identified populations of lymph node stromal cells (LNSCs) that retain antigen for 14 days post-vaccination and gene expression patterns associated with antigen archiving^11^. The ability of a LEC to archive antigen could be influenced by two separate processes: 1) antigen uptake, which likely occurs through endocytic pathways, and 2) antigen retention, which would be influenced by the cellular compartment where the archived antigen is stored. However, our previous studies^11^, lacked the temporal resolution to differentiate between these two processes. Here we sought to evaluate the dynamics of antigen uptake and archiving by extending the timeframe to encompass a three week and six week post-immunization time-course.

We immunized mice with ova-psDNA + VV to induce robust LN expansion and antigen archiving^8, 11^ and performed scRNA-seq on CD45-cell populations from the draining lymph nodes at 21 days and 42 days post-vaccination. In a separate cohort of mice immunized at the same time, we sorted LN myeloid cell populations, which were enriched for CD11c+, CD11c-CD11b+, B220+ and all other cells. Sorted populations were mixed a 4:4:1:1 ratio, respectively^11^. We then compared antigen levels across a time-course using our published scRNA-seq data collected 2 and 14 days post-immunization^11^. To accurately compare cell populations across the four time points, we integrated the scRNA-seq data for each time point and re-annotated the cell types present in each sample using a previously published automated approach^21^ (Figure S1A). To compare levels of antigen across our datasets we calculated background-corrected antigen scores (Ag-score) for each cell, normalizing counts by the total antigen counts for the sample, subtracting the estimated background signal based on counts present in B and T cells, and log-transforming (see methods). In support of our previous work, we detected the highest antigen levels within lymphatic endothelial cells (LECs), followed by lower levels in neutrophils, blood endothelial cells (BECs), fibroblasts, monocytes, and dendritic cells (DCs) (Figure 1B, S2A-B). Levels of archived antigen were highest 2 days post-vaccination (Figure 1B). In all cell types detected, except LECs, antigen levels dropped sharply by day 14 and were largely undetectable by day 42. LECs maintained high levels of antigen throughout the 21 day time-point, however by day 42 antigen levels were substantially reduced, though still higher than all other cell populations. These results support our previous work showing that LECs are the predominant cell population that acquires and archives antigens in the draining lymph node^11^.

**Figure 1.**
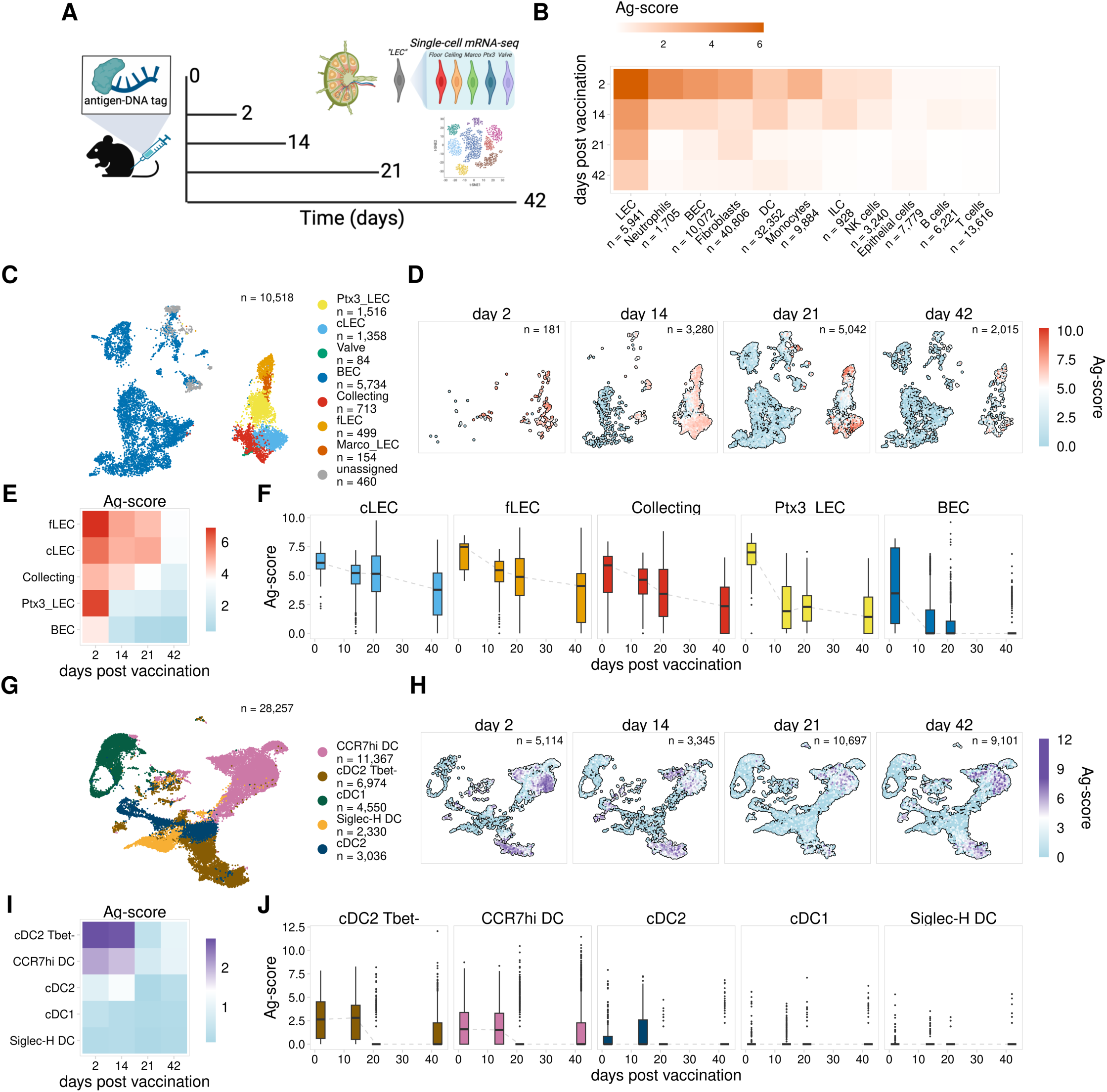
Antigen persists in discrete cell populations within the lymph node A. Experimental design B. Mean Ag-score is shown for each cell type for 2, 14, 21, and 42 days post vaccination. C. UMAP projection shows LEC subsets. D. UMAP projections show Ag-scores for LEC subsets for each timepoint. E. Mean Ag-score is shown for LEC subsets for each timepoint. F. Ag-scores are shown for each timepoint for each LEC subset. G. UMAP projection shows DC subsets. H. UMAP projections show Ag-scores for DC subsets for each timepoint. I. Mean Ag-score is shown for DC subsets for each timepoint.Ag-scores are shown for each timepoint for each DC subset. J. Ag-scores are shown for each timepoint for each DC subset.

To investigate the dynamics of antigen acquisition and archiving across different LEC subsets, we next annotated different endothelial cell populations using previously published data^11, 14^ (Figure S1B, C). Using this approach, we identified subsets of collecting and valve LECs that form the afferent and efferent collecting vessels of the lymph node (collecting LECs) (Figure 1C, S1B-C), as well as floor (fLEC) and ceiling (cLEC) of the subcapsular sinus (SCS), which express the adhesion molecule Madcam1 and the scavenger receptor Ackr4, respectively (Figure 1C, S1B-C). SCS LECs are the first LEC populations that encounter lymph-borne antigens after entry into the lymph node and we previously showed that these populations have the highest levels of antigen 14 days post-immunization^11^. We also identified LEC subsets from the medullary and paracortical sinuses, which are distinguished by the expression of the scavenger receptors MARCO (Marco LECs) and Ptx3 (Ptx3 LECs)^11, 14, 16, 17^ (Figure 1C, S1B-C). Analysis of antigen levels over time for each LEC population revealed that at 2 days post-immunization, LECs show universally high levels of tagged antigen (Figure 1D-F, S2C). These findings indicate that immediately following a protein subunit immunization, antigen is dispersed throughout the LN, including in Ptx3 LECs, which represent one of the last populations to encounter antigen based on the location of Ptx3 LECs identified by others^14^. However, antigen levels at later times after immunization showed significant differences in antigen archiving ability. We detected high antigen levels in cLECs, fLECs, and collecting LECs through 21 days post-immunization (Figure 1D-F), while antigen levels drop significantly by day 14 in Ptx3 LECs (Figure 1D-F, S2C). These results indicate that differences in antigen archiving are not due to LEC localization within the lymph node or efficiency of antigen uptake, as we observe high amounts of antigen within all LEC populations 2 days after immunization, and instead suggest that the dynamics of antigen archiving are a key determinant of overall antigen levels.

The protective capacity conferred by antigen archiving is dependent on antigen exchange with migratory DCs, which present archived antigen peptides to memory CD8 T cells^8,9^. To explore the dynamics of antigen exchange between LECs and different DC subsets, we integrated the DC datasets for each time point and annotated specific DC populations using previously published data^11, 22, 23^, identifying conventional (c) DC1 and DC2 populations along with CCR7hi migratory cDCs and Siglec-H-expressing plasmacytoid DCs (Figure 1G, S1D-E). Comparison of antigen levels between these populations showed cDC2 Tbet- and CCR7hi DCs had the highest antigen levels 2 days post-vaccination, with other subsets having minimal levels at any time point (Figure 1H-J, S2D). The CCR7hi and Tbet-DC subsets both showed consistent levels of antigen through day 14, with levels dropping sharply by day 21. These findings are consistent with other reports demonstrating the length of time DCs remain viable after activation^24^. Therefore, we suggest that any antigen remaining within the DC populations beyond 2-3 weeks is likely a result of DC acquisition of antigen from other cell types. Consistent with our published data that migratory DCs are required for antigen exchange, it is likely the antigen positive DCs remaining have acquired antigen from the only cell type with antigen at these late time-points, LECs.

### Identification of gene signatures associated with antigen archiving by LECs

Comparison of antigen levels throughout the time course revealed significant differences in antigen archiving between LEC populations. We next identified gene expression signatures that might influence the ability of different LEC subsets to archive antigen. We previously identified genes associated with caveolin-mediated endocytosis that are expressed more strongly in LECs with higher levels of archived antigen at day 14^11^. We sought to leverage our new scRNA-seq data to expand on this idea and further characterize gene expression programs underlying antigen archiving. We first identified LECs with high levels of antigen (Ag-high) at the day 14 time point by performing k-means clustering separately for each LEC subset using the calculated antigen scores (Figure 2A). To ensure consistent classification of Ag-high cells across the time points, we used the day 14 metrics to classify antigen-high (Ag-high) cells in all other time points.

**Figure 2.**
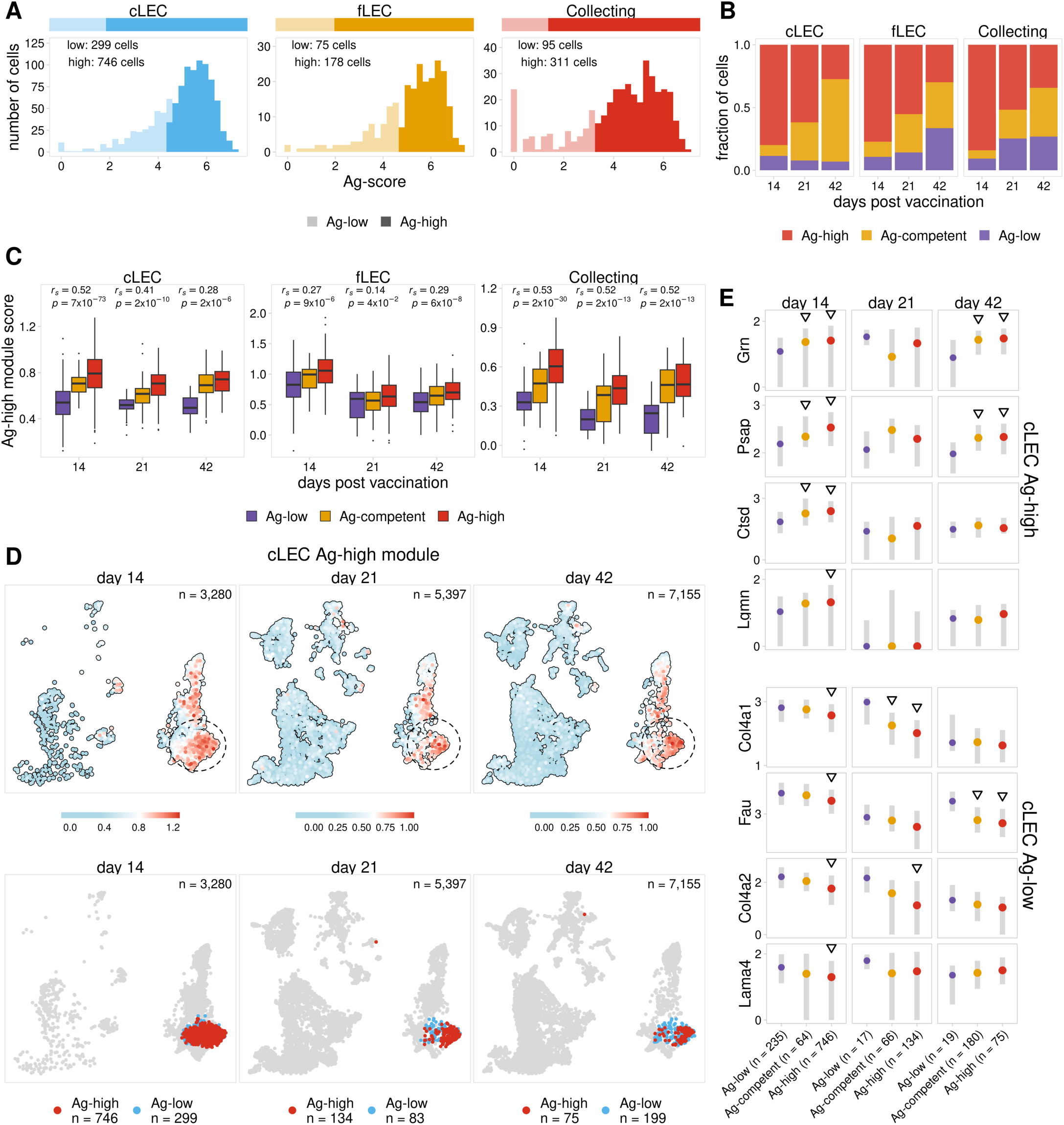
Identification of gene signatures associated with antigen archiving by LECs A. Ag-score is shown for day 14 Ag-high and -low cLECs, fLECs, and collecting LECs. B. The fraction of cells predicted to be Ag-competent is shown for each LEC subset 14, 21, and 42 days post vaccination. C. Ag-high module scores are shown for Ag-low, Ag-high, and predicted Ag-competent LECs. The Spearman correlation between Ag classes and Ag-high module score is shown for each timepoint. One-sided p values were calculated and adjusted using Benjamini-Hochberg correction. D. UMAP projections show cLEC Ag-high module scores for each timepoint. E. The expression of select genes from the Ag-high (top four) and Ag-low (bottom four) gene modules is shown for cLECs. Triangles indicate the gene is differentially expressed when compared to Ag-low cells. P values were calculated using a one-sided Wilcoxon rank sum test with Benjamini-Hochberg correction.

To investigate gene expression patterns underlying antigen retention, we next identified genes that were differentially expressed between Ag-high and -low populations. Using these genes, we trained separate random forest classifiers using day 14 cLECs, fLECs, and collecting LECs. Using these models, we identified genes most predictive of whether an LEC from one of these subsets would acquire high or low amounts of antigen. We then grouped these genes into distinct Ag-high and Ag-low gene programs based on whether the gene was up- or down-regulated in Ag-high cells, respectively. To characterize cellular processes potentially influencing antigen archiving, we identified gene ontology terms for the Ag-high gene program for each LEC subset. From this analysis, we find that the Ag-high programs overlap with ontology terms suggestive of increased endosomal-lysosomal function (Figure S3A), and this is most significant for cLECs and fLECs. This includes lysosome factors with broad roles in protein homeostasis (Ctsd^25^, Grn^26^, Lgmn^27^, Prcp^28^), lipid metabolism (Psap^29^), and lysosome acidification (Atp6v1a^30^). In addition, expression of the clathrin adaptor protein Dab2^31^ may indicate increased endocytic activity. To further investigate how the Ag-high gene programs could affect antigen archiving, we next determined whether these genes were induced in response to an immune stimulus. We compared each time point after immunization with previously published scRNA-seq data for sorted LNSCs from naive mice^17^. We found that this program is robustly expressed in LECs from naive mice and expression persists 42 days post-vaccination when most archived antigen has been released or degraded (Figure S3C-D). Taken together, these results suggest that the identified gene programs are uniquely expressed by most lymphatic endothelial cells regardless of inflammatory state, and not induced in response to immune stimulus. Why LECs have increased expression of genes associated with endo-lysosomal trafficking is unknown, however it seems likely that LECs must degrade antigens over time. The high levels of antigen acquired by LECs are therefore likely to contribute to the length of antigen duration, which is consistent with prior work^10^.

Continued expression of the Ag-high gene program after 21 and 42 days post immunization (Figure S3C) suggests there are still LECs that are competent to archive antigen even though most antigen has been degraded or released by these time points. To further investigate these potential archiving-competent cells, we attempted to identify LEC populations that would be predicted to archive antigen at the 21 and 42 day time points, based on the classification models trained using the day 14 time point. From this analysis we classified cells into three groups: 1) Ag-low has low amounts of archived antigen based on quantification of antigen scores, 2) Ag-competent has low amounts of antigen but is predicted to have high antigen levels by our classification models, and 3) Ag-high has high levels of antigen. We find that at day 21 and 42 most cLECs, fLECs, and collecting LECs that have low levels of antigen are predicted to be Ag-competent (Figure 2B, S3E). To assess the accuracy of these predictions, we compared expression of the Ag-high and Ag-low gene modules derived from our day 14 classification models. As expected, we found that the predicted Ag-competent cells have higher expression of the Ag-high module and reduced expression of the Ag-low module when compared with Ag-low cells (Figure 2C, S3F). Furthermore, by comparing Ag-low, Ag-competent, and Ag-high cells, we found a notable correlation between these classes and the expression of the Ag-high and -low gene programs (Figure 2C, S3F). We saw that LECs that retain high amounts of antigen at the 21 and 42 day time points generally show the strongest and weakest expression of the Ag-high and -low programs, respectively (Figure 2C-E, S3F-H). To summarize, we were able to identify gene signatures (Figure S3A-B) expressed in LECs from naive mice (Figure S3C-D) that are predictive of the capacity of a cell to archive antigen (Figure 2B, S3E, Table S1) and correlate with the duration of antigen retention (Figure 2C, S3F).

### Antigen archiving by LECs is enhanced by sequential immunization

We have previously shown that a subsequent immune challenge results in the presentation of previously archived antigens, which in turn boosts protective immunity^14^. To expand on these results, we next investigated how the dynamics of antigen archiving are impacted by sequential immunizations administered within the archiving time frame. We compared mice from three vaccination groups, A) received one vaccination 42 days prior to harvest, B) received one vaccination 21 days prior to harvest, and C) received both vaccinations, 42 and 21 days prior to harvest (Figure 3A). For each group we immunized WT mice as above with ova-psDNA+VV subcutaneously, harvested draining popliteal lymph nodes, and performed scRNA-seq on LN populations as described above. To separately quantify archived antigen originating from the 21 day (red syringe) and 42 day (blue syringe) time points, the psDNA tags for each immunization contained unique barcodes (Figure 3A).

**Figure 3.**
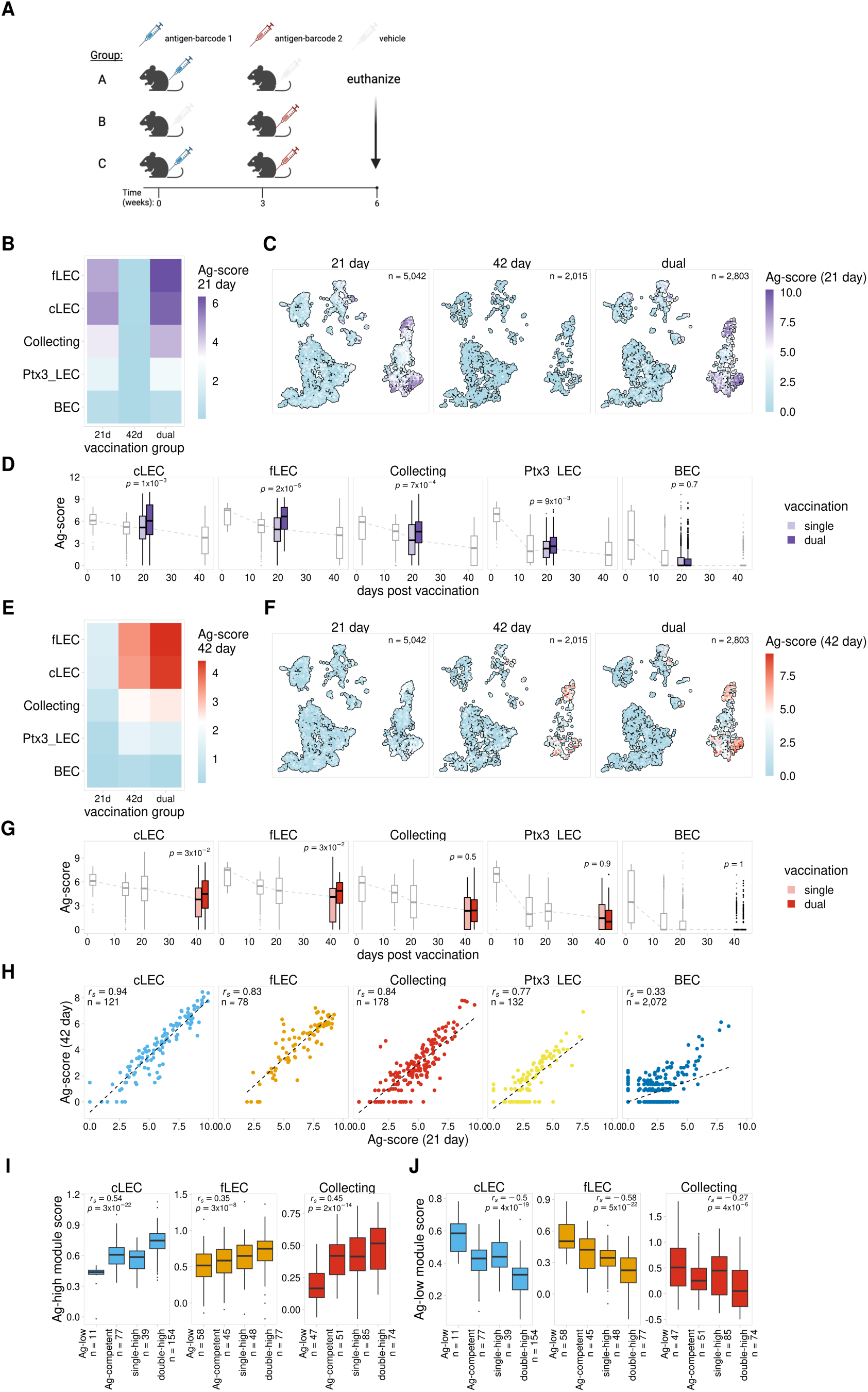
Antigen archiving by LECs is enhanced by sequential immunizations A. Experimental design B. Mean 21 day Ag-score is shown for LECs from mice that received a single vaccination (21 day or 42 day) or dual vaccination (21 day and 42 day). C. UMAP projections show 21 day Ag-scores for LEC subsets for single and dual vaccinations. D. Prior vaccination enhances antigen archiving. Ag-score is shown for single and dual vaccinations for the 21 day timepoint for each LEC subset. Other timepoints are shown in grey. P values were calculated using a one-sided Wilcoxon rank sum test with Benjamini-Hochberg correction. E. Mean 42 day Ag-score is shown as described in A. F. UMAP projections show 42 day Ag-scores for LEC subsets as described in B. G. Successive vaccinations enhances retention of previously archived antigen. Ag-score is shown for the 42 day timepoint as described in C. H. 21 day and 42 day Ag-scores are compared for LEC subsets. I. Ag-high module scores described in Figure 2 are shown for LECs that archived antigens from both vaccinations (double-high), from only one vaccination (single-high), have low levels of both antigens, but are predicted to be Ag-competent, or have low levels of both antigens (Ag-low). Module scores are shown for the corresponding LEC subset. P values were calculated using a one-sided Wilcoxon rank sum test with Benjamini-Hochberg correction. J. Ag-low module scores are shown for LEC subsets as described in H.

To first investigate how previously archived antigen affects archiving during a future immune challenge, we compared levels of the 21 day antigen for LECs from group B, which only received the second immunization (21 day), and group C, which received both immunizations. Consistent with our previous results (Figure 1), the LEC subsets with the highest levels of antigen in the dual vaccinated mice (group C) are cLECs, fLECs, and collecting LECs (Figure 3B). As expected, we also observed minimal background levels of the 21 day antigen for mice that did not receive the 21 day immunization (Figure 3B-C). Comparison of the 21 day antigen levels for single vs dual vaccinated mice revealed elevated levels of archived antigen in mice that received both the 42 and 21 day immunizations (Figure 3B-D). This suggests that LECs that previously archived antigen have enhanced uptake and/or retention of antigens from a future immunization. Whether this function of increased antigen uptake is permanent and the result of an altered transcriptional program caused by antigen acquisition or inflammation is still unclear.

Using this dual vaccination model, we next tested the inverse scenario and asked how successive immunizations affect retention of previously archived antigen. To do this we compared levels of the 42 day antigen for mice that only received the first vaccination (group A) vs mice that received both vaccinations (group C) (Figure 3A). Similar to the results for the 21 day antigen, we saw significantly higher levels of the 42 day antigen in dual vaccinated mice, suggesting prolonged retention of the previously archived antigen (Figure 3E-G, S4B). Taken together, these results indicate that sequential vaccinations promote an overall increase in antigen archiving by LECs.

To further investigate the dynamics of antigen uptake and retention, we next asked whether increased antigen levels observed in dual vaccinated mice was due to distinct groups of LECs acquiring each antigen. Evaluation of LECs from the 42 day time point indicated a substantial fraction of cells with low levels of antigen that expressed a gene program predictive of increased antigen archiving ability (Ag-competent, Figure 2B). This represents a cell population that could be well suited to take up new antigen during a future immune challenge. From this observation, we predicted that in dual vaccinated mice we would observe two distinct groups of Ag-high cells, one primarily containing antigen from the first vaccination (42 day) and a separate group containing antigen from the second vaccination (21 day). To test this model, we compared levels of each tagged antigen in the dual vaccinated mice. Contrary to our prediction, we did not observe distinct groups of LECs that exclusively archived the different antigens, but instead found a strong correlation between levels of the tagged antigens for each LEC subset, with cells containing high levels of the first antigen (42 day) also showing high levels of the second antigen (21 day) (Figure 3G, S4D). We confirmed these findings *in vivo* using fluorescently labeled ovalbumin with polyI:C/aCD40, and similarly found that the same LECs from draining LNs of mice immunized 1 week apart were positive for both antigens delivered sequentially (Figure S5), indicating there are populations of cells within each LEC subset that are particularly well suited to archive antigen and will acquire antigens from repeated immune challenges.

Finally, we next asked whether there were distinct gene expression signatures associated with cells that archived multiple antigens in the dual immunized mice. We analyzed expression of the previously identified Ag-high and Ag-low gene modules (Figure 2) to determine if these programs might also influence the ability of a LEC to archive antigens from multiple vaccinations. We identified LECs from the dual immunized mice that had high levels of each antigen using the antigen score cutoffs derived from the day 14 time point (Figure 2A) and predicted Ag-competent cells using our previously described classification models (Figure 2, S3). We then identified LECs with high levels of both antigens (double-high), high levels of only one antigen (single-high), low levels of both antigens but predicted to be Ag-competent (Ag-competent), and low levels of both antigens (Ag-low). Comparison of these groups reveals a significant correlation with expression of the Ag-high and Ag-low gene programs (Figure 3I-J, S4E-F). We found that the double-high cells, which are the most proficient at archiving, had the highest expression of the Ag-high gene program and the lowest expression of the Ag-low gene program for each LEC subset (Figure 3I-J, S4E-F). These findings suggest that we have identified at least some of the gene programs that promote antigen archiving during an initial immunization and that these programs also function to promote archiving during sequential immune challenges.

### Antigen exchange with DCs correlates with levels of antigen archiving

We have previously shown that to enhance protective immunity, LECs exchange archived antigen with migratory DCs, which present to memory CD8 T cells^9, 10^. To further investigate antigen exchange between LECs and DCs, we assessed the spatial localization of antigen in the draining lymph node using the GeoMx DSP spatial transcriptomics platform. We repeated our dual vaccination experiment and immunized mice with ova-psDNA for 21 days, 42 days, or sequentially 21 days apart using ova-psDNA conjugates containing unique barcodes with VV (Figure 3A). We excised draining popliteal LNs at each time point, fixed with formalin, and embedded into paraffin. Tissue sections were cut and analyzed with the GeoMx DSP platform using probes for the mouse transcriptome, custom probes recognizing the 21 day and 42 day antigen barcodes, and negative control probes for normalization. We obtained data for a total of 15 individual lymph nodes divided across two slides (Figure S6A). Distinct regions (composed of >500 cells on average) within each lymph node were selected for analysis, including the subcapsular sinus, medulla, and cortex (Figure 4A, S6B). The GeoMx platform allows each selected region to be further segmented based on fluorescent staining. To distinguish between different cell populations, each region was segmented based on antibody staining for Lyve1 (LECs), CD11c (DCs), and CD4 (T cells) (Figure 4B).

**Figure 4.**
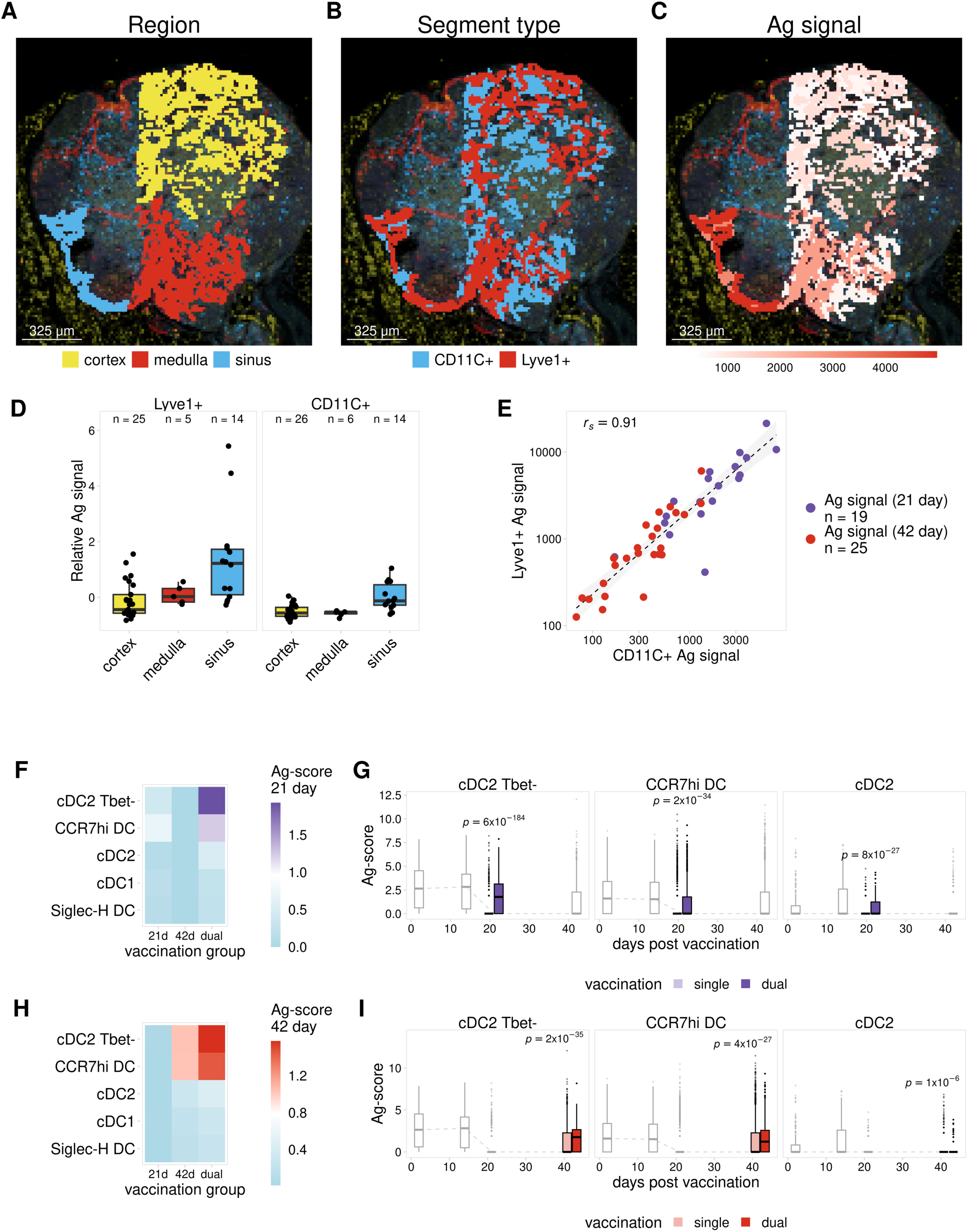
Antigen exchange with DCs correlates with levels of antigen archiving A. Annotated regions used for antigen quantification are shown for a representative lymph node. B. Segment type is shown as described in A. C. Antigen signal is shown as described in A. D. Relative antigen tag signal is shown for each sample and each region segmented based on Lyve1 (LECs) and CD11c (DCs). Normalized 21 day antigen signal was scaled across all regions from 21 day mice and dual immunized mice, 42 day antigen signal was scaled across all regions from 42 day mice and dual immunized mice. The number of plotted segments is shown above each boxplot. E. Normalized antigen tag signal is shown for regions that included both Lyve1+ and CD11c+ segments. 21 day antigen signal is shown for 21 day and dual immunized mice, 42 day signal is shown for 42 day and dual immunized mice. The Spearman correlation coefficient is shown. F. Mean 21 day Ag-score is shown for DCs from mice that received a single vaccination (21 day or 42 day) or dual vaccination (21 day and 42 day). G. Ag-score is shown for single and dual vaccinations for the 21 day timepoint for DC subsets with the highest Ag-scores. Other timepoints are shown in grey. P values were calculated using a one-sided Wilcoxon rank sum test with Benjamini-Hochberg correction. H. Mean 42 day Ag-score is shown as described in F. I. Ag-score is shown for single and dual vaccinations for the 42 day timepoint as described in G.

To identify regions with the highest levels of antigen, we next compared the normalized antigen signal for probes targeting the 21 day and 42 day antigens. As we previously showed with our scRNA-seq experiments (Figure 1), we found that antigen signal is higher at the 21 day time point and drops substantially by day 42 (Figure S6C-D). To compare antigen signals for each region, we pooled regions from the single and dual immunized mice. We then scaled the normalized signal separately for each antigen to allow the 21 and 42 day antigen signals to be compared together. As expected, we found that antigen levels were highest within Lyve1+ segments corresponding to LECs (Figure 4B-D). In addition, antigen levels for both Lyve1+ and CD11c+ segments are highest within subcapsular sinus regions (Figure 4D, S6C). This supports our previous scRNA-seq results showing that LEC populations making up the ceiling and floor of the subcapsular sinus (cLECs and fLECs) contain the highest levels of archived antigen (Figure 1D-F)^11^. We next asked whether the localization of DCs to different regions of the lymph node influenced the amount of antigen acquisition by DCs. Similar to the Lyve1+ segments, we found that CD11c+ segments in the subcapsular sinus had higher levels of antigen (Figure 4D, S6C- D). In addition, there was a strong correlation between antigen signals for Lyve1+ and CD11c+ segments within each region, and regions with higher antigen levels in LECs were also higher for antigen in DCs. This suggests that the amount of antigen exchanged with DCs is directly dependent on the amount of antigen archived by proximal LEC populations where SCS LECs archive the most antigen and therefore DCs located near the SCS or migrating through the SCS also have the highest amount of antigen.

We next asked if there was increased antigen uptake by DCs in dual vaccinated mice, which showed higher levels of archived antigen in LEC populations (Figure 3B-G). Due to the low number of regions obtained for dual immunized mice (Figure S6A-C), we were not able to compare antigen signals using the GeoMx platform. Instead, we used our scRNA-seq data to analyze DC populations from single- and dual-immunized mice (Figure 3A). In support of our previous results (Figure 1H-J), we found that CCR7hi and cDC2 Tbet-populations had the highest antigen levels in the dual-vaccinated samples. In addition, similar to LECs (Figure 3B-G, S4A-C), we found that DCs from dual vaccinated mice had significantly higher antigen levels than mice that only received one of the immunizations (Figure 4F-I, S4A,C). These results in conjunction with our spatial transcriptomics data (Figure 4A-E), support a model wherein successive vaccinations enhances antigen uptake and archiving, which in turn results in increased exchange of antigen with proximal DC populations.

### Antigen archiving is impaired during CHIKV infection

By training classification models for the main LEC subsets that archive antigen, we identified cells that are predicted to be archiving-competent (Ag-competent) based on the expression of distinct gene programs (Figure 2). To further test the utility of these models, we asked if our models could predict differences in the antigen archiving capability of LECs between naive mice and mice infected with CHIKV, which impairs LN antigen acquisition^16^. We used previously published scRNA-seq data of murine LNSCs 24 hours after infection with the arthritogenic alphavirus CHIKV^16, 17^. CHIKV infection in mice disrupts the structure and function of the draining lymph node^17, 20^ and induces a proinflammatory and antiviral response in LECs^16^. Importantly, by 72 hours post-infection with WT CHIKV, antigen acquisition by LN LECs is impaired^16^. Using scRNA-seq data for lymph node stromal cells at 24 hours post CHIKV infection (Figure 5A), we asked if we could use our antigen archiving classification models to demonstrate whether CHIKV infection would also impede antigen archiving at 24 hours post infection.

**Figure 5.**
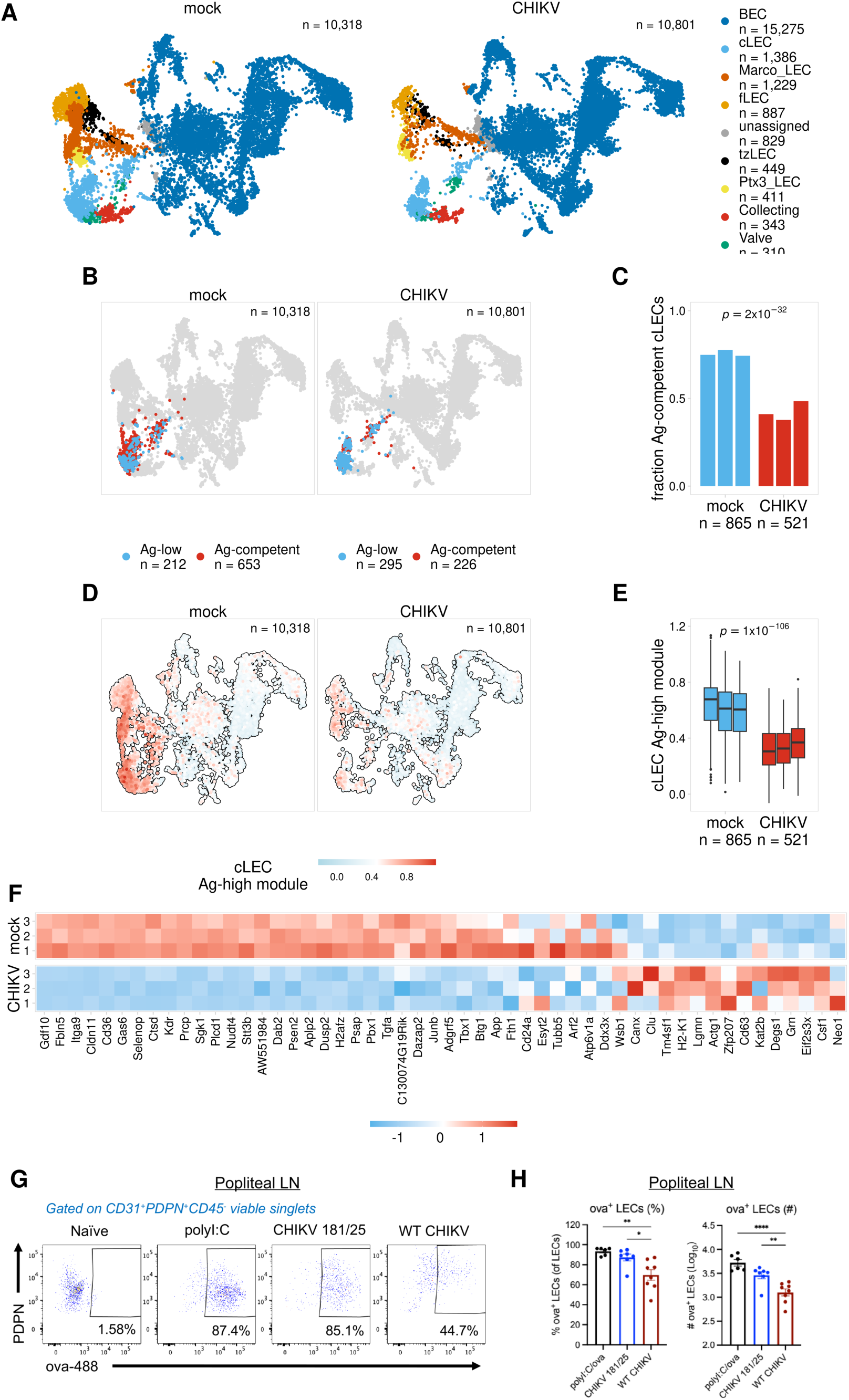
Antigen archiving is impaired during CHIKV infections A. UMAP projections show LEC subsets for mock and CHIKV-infected mice. B. UMAP projections show predicted Ag-competent cLECs for mock and CHIKV-infected mice. C. The fraction of predicted Ag-competent cLECs is shown for mock and CHIKV-infected mice for each biological replicate. P values were calculated using Fisher’s exact test with Benjamini-Hochberg correction. D. UMAP projections show cLEC Ag-high module scores for mock and CHIKV-infected mice. E. cLEC Ag-high module scores are shown for mock and CHIKV-infected mice for each biological replicate. P values were calculated using a two-sided Wilcoxon rank sum test with Benjamini-Hochberg correction. F. Expression of the cLEC Ag-high gene module is shown for mock and CHIKV-infected mice for cLECs from each biological replicate. G. Representative flow cytometry plots showing ova+ LECs H. The percentage and total number of ova+ LECs

The scRNA-seq data for CHIKV-infected mice includes similar subsets of LECs, including cLECs, fLECs, and collecting LECs (Figure 5A). To evaluate antigen archiving during CHIKV infection, we predicted archiving ability for cLECs, fLECs, and collecting LECs from mock- and CHIKV-infected samples using the classification models previously trained for each of these subsets (Figure 5B-C, S7C). To first assess the accuracy of these predictions, we compared expression of the Ag-high and Ag-low gene modules derived from each model (Figure 2). As expected, there was overall higher expression of the Ag-high gene module and lower expression of the Ag-low gene module for LECs predicted to be archiving-competent (Ag-competent) (Figure S6A). We next compared the fraction of Ag-competent cells for the mock-and CHIKV-infected samples. We found that cLECs had the largest change, with 75% of cells predicted to archive antigen in mock-infected mice and only 43% of cells predicted to archive antigen in CHIKV-infected mice (Figure 5B-C, S7C). Comparison of expression differences for the Ag-high and Ag-low gene modules for mock- and CHIKV-infected mice further supported these findings. For cLECs we found a significant reduction in expression of the Ag-high module and an increase in expression of the Ag-low module during CHIKV infection (Figure 5D-F, S7B). However, fLECs and collecting LECs both had less pronounced changes in the fraction of Ag-competent cells and in the expression of these modules. fLECs displayed a reduction in both Ag-high and -low modules, and collecting LECs only showed subtle differences in the expression of each. These results suggest a reduction in antigen archiving ability at 24 hours post infection, and this effect appears most prominent within the cLEC population.

To experimentally test the accuracy of these predictions, we infected mice in the footpad with WT CHIKV or CHIKV 181/25, which does not impair antigen acquisition by LN LECs^16^. After 24 hours, mice were immunized with 10 μg ova-488 in both calf muscles (20 μg/mouse total), and ova^+^ LECs were evaluated in the dLN 2 days later. As a positive control, mock-infected mice were immunized with 10 μg ova-488 and 5 μg polyI:C in both calf muscles. Consistent with our previous work, the percentage and number of ova^+^ LECs was similar between CHIKV 181/25-infected and polyI:C-treated mice (Figure 5G-H). However, there was a significant reduction in the percentage and number of ova^+^ LECs in WT CHIKV-infected mice compared with CHIKV 181/25-infected mice (Figure 5G-H). These data indicate that antigen acquisition, the first step in antigen archiving, is impaired during WT CHIKV infection as early as 24 hours after inoculation, prior to the alterations in LN LEC composition and LN cellular organization previously shown during WT CHIKV infection^16, 20^. Thus, these findings support the accuracy of our predictive models using gene expression patterns to inform LEC antigen archiving capability, and illustrates the utility of these tools to assess antigen archiving in systems where inclusion of antigen-psDNA reagents are not feasible.

## Discussion

We have defined a gene signature associated with LEC antigen archiving using an antigen conjugated to a DNA tag for use with single cell RNA sequencing. LECs acquire and archive foreign antigens beyond the peak of the primary immune response^8–11^. We previously demonstrated that antigen can be detected using a psDNA tag conjugated to a protein antigen for up to 14 days post-immunization, using single cell sequencing^11^. We further extended this timeline to 21 and 42 days post-immunization to assess the dynamics of antigen archiving. Corroborating our published data, we found that antigens persist predominantly in LEC subsets, such as cLEC, fLEC, and collecting LEC, at late time points following both single and dual vaccinations (Figure 1F, 3G). Interestingly, Ptx3 LECs capture antigens at day 2, but by day 14 antigen levels decline in these cells. One potential explanation for the loss of antigens within the Ptx3 LEC subset may be the lack of expression of *Cd274* (PDL1). PDL1 is an important regulator that determines LEC division and survival where loss of PD-L1 leads to increased LEC division^32^. Indeed, the LECs that undergo division during LN expansion following immunization are also the LECs that undergo apoptosis^32^. Thus, Ptx3 LECs, which have low expression of *CD274*^14^, likely undergo apoptosis and lose antigens earlier than the other LEC subsets during LN contraction.

We developed machine learning models to better predict cells capable of antigen archiving. The training of the classification model was based on the day 14 timepoint because it was an intermediate timepoint with robust antigen high and low populations. As a proof of concept, these models were able to predict the decrease in Ag-competent cells following CHIKV infection. Extensive data^16^ demonstrate how CHIKV infection disrupts the organization of the LN through interactions with LNSCs. We further tested the accuracy of these predictions by assessing the ability of LECs to archive fluorescent antigens in CHIKV-infected mice (Figure 5). The novelty in using these classification models is the ability to apply the classifications to already available scRNAseq datasets from different infections or immunizations and assess antigen competency. This could be particularly important to identify which infections or inflammatory stimuli may prohibit or reduce antigen archiving and may inform studies regarding when or which area to vaccinate. Upon expanding on our published study using antigen-DNA conjugates as a molecular tracking system, we demonstrated that the incorporation of two different antigen tags can accurately model the kinetics of antigen signals in different cell types using scRNA-seq. Furthermore, the development of machine learning models to assess antigen archiving presents a novel approach for future applications.

Genes associated with endosomal-lysosomal function were captured as part of this antigen archiving gene signature. Lysosomal-associated genes in the Ag-high module (Figure S3A) could result from the large amount of antigens within the LECs. It seems likely that LECs trap foreign antigens as they pass through the LN similar to how LECs capture viral particles to prevent viral dissemination^17^. Ultimately the LECs must degrade or release antigens to passerby cells, such as DCs. Antigen exchange between LECs and DCs is likely a by-product of the process of antigen archiving and maintained as small amounts of DC antigen presentation over a limited time period are likely an evolutionary benefit. Indeed, we have previously demonstrated an increased capacity for DC presentation of archived antigens during an unrelated infection^8^. During infection, the increased activation state of the DCs that acquire both antigens from the infection and antigens from LECs do appear to lead to improved memory T cell responses and increased effector function^8^. Alternatively, LECs could acquire antigens via different endocytic pathways. As an example, we and others have demonstrated that LEC endocytosis of proteins occurs via caveolin and clathrin mediated endocytosis *in vivo*^11^ and *in vitro*^33^. Caveosomes maintain a neutral pH and cargo can remain in caveosomes until transcytosis/recycling or lysosomal degradation via RAB5 dependent fusion with the early endosome^34^. It is possible that caveolin mediated endocytosis is one mechanism of antigen storage while clathrin mediated endocytosis targets excess antigen for degradation. The data presented here suggest that specific LECs are “pre-coded” to be competent in antigen archiving, particularly when our classification models were applied to naïve mice (Figure 5). These models predict that certain LECs within naïve mice are predisposed to retain the antigen-archiving gene signature (Figure S3C-D).

As LECs do not directly present archived antigens to CD8 T cells, there is a need for antigen exchange from LEC to migratory DC for the presentation of archived antigen-derived peptides^8, 9, 11^. Indeed, antigen-bearing LECs interact, particularly those that express ICAM1, and exchange antigen with DCs^9^. Building upon the live imaging of LEC-DC interactions to exchange archived antigens, here we visualized the spatial localization of antigen bearing cells in the LN using the Nanostring GeoMx platform. We confirmed by scRNA-seq that antigen signals within LEC and DC subsets correlated with our spatial data. We found that CCR7hi DC and cDC2 T-bet DC had antigen levels peaking at 14 days post-vaccination and that antigen levels dramatically decreased by day 21 post-vaccination (Figure 1J). Interestingly, when we further extended this time course to 42 days, we still detected antigens in cDC2 Tbet- and CCR7hi DC populations (Figure 1J). While it is unlikely that DCs are still acquiring and processing antigens from the initial immunization several weeks earlier, this finding supports the hypothesis that LECs are slowly releasing antigens over time through either direct LEC-DC interactions or via LEC apoptosis.

In dual-vaccinated mice, cLECs, fLECs, and collecting LECs have enhanced antigen acquisition and retention compared to mice receiving a single vaccination (Figure 3D, G). The data we provide outline: *i.* that LECs are capable of acquiring multiple antigens and *ii*. that LECs that acquired the first antigen are even better at acquiring a second antigen. We found that the gene signature for LECs during one immunization is similar to the gene signature of LECs with multiple archived antigens. This suggests there are LECs that are more susceptible to being reprogrammed for antigen acquisition and archiving (Ag-competent). It is possible that LNSCs, particularly LECs, undergo epigenetic reprogramming during an initial antigen encounter that provides increased responsiveness in regards to antigen acquisition. Trained immunity is thought to be mediated by epigenetic changes, alterations in gene expression, and metabolic reprogramming within innate immune cells. These changes lead to a heightened state of readiness and elimination of pathogens upon subsequent exposure^35, 36^. Interestingly, this training is not limited to innate immune cells as one study showed that IFNb stimulation contributes to increased and quicker induction of interferon-stimulated genes in mouse embryonic fibroblasts^37^. Upon restimulation, the rapid induction of ISG was attributed to increased recruitment of polymerase II to ISG loci and this was also associated with H3.3 and H3K36ME3 marks on the chromatin of ISG genes^37^. Thus, it is possible that LECs may undergo epigenetic modifications upon antigen acquisition to promote subsequent antigen uptake and retention.

We found higher antigen levels in both LEC subsets and DC subsets, CCR7hi DC and cDC2 Tbet-, upon a second antigen exposure compared to single-immunized mice (Supp. Figure A, C). This supports our published data demonstrating that migratory DCs^9^ and cDC1s are required for cross-presentation of archived antigens during the time frame of LEC apoptosis^8^. cDC1 cross-presentation of cell-associated antigens is partly attributed to their ability to internalize apoptotic cells or cell debris^38–41^, thus when mice receive a second vaccination, it is possible that some of LECs undergo apoptosis. LEC apoptosis causes the release of archived antigens which DCs can acquire. Alternatively, DCs may have higher levels of antigen after a second exposure for the same reason as LECs. Myeloid cell populations do undergo innate training and thus perhaps DCs are also capable of increasing antigen acquisition in addition to increased cytokine release as has been described^35, 42, 43^.

A caveat to our studies is the inadequate number of regions we could evaluate in doubly immunized mice to assess the two different antigen tags spatially in the LN. However, because the GeoMx spatial transcriptomic platform lacks the depth of scRNA-seq, we coupled the GeoMx platform with our single-cell RNA-seq datasets. Here, we found that Lyve1+ and CD11c+ sections of the LN had the highest level of antigens at the subcapsular sinus of the LN. These data support our finding that sequential vaccinations enhanced LEC antigen archiving and antigen exchange. Together this manuscript provides confirmatory evidence of our prior publications using transcriptional profiling and antigen DNA tags. To build upon these data we provide a unique resource to assay antigen archiving capacity by developing an antigen archiving gene signature and a unique antigen competency model to predict outcomes from other datasets. These models could also prove to be appropriately suited for modeling human antigen archiving. Indeed, preliminary analysis suggests a similar gene expression program is available in human LEC populations from existing datasets^44^.

## Materials and Methods

### Mice

5-6 week-old mice were purchased from Charles River or Jackson Laboratory, unless otherwise stated, bred and housed in the University of Colorado Anschutz Medical Campus Animal Barrier Facility. Wild type were all bred on a C57BL/6 background. All animal procedures were approved by the Institutional Animal Care and Use Committee at the University of Colorado.

### Immunizations and Infections

6–8-week-old C57BL/6 (CD45.2) mice were immunized with 1E4 plaque-forming units (PFU) of Vaccinia Western Reserve or 5 µg of poly I:C (Invivogen) with or without 5 µg of anti-CD40 (FGK4.5, BioXcell) and 20 µg of ova-psDNA or ova in 50 μL volume per footpad injection. Endotoxin levels were quantified using the Pierce Limulus Amebocyte Lysate Chromogenic Endotoxin Quantitation kit (Thermo Scientific) to be less than 0.5 EU/mg for either ova or ova conjugated to psDNA. For CHIKV infections, 4 week old C57BL/6 mice were inoculated with 10^3^ PFU of CHIKV 181/25 or CHIKV AF15561 (WT CHIKV) in a 10 μL volume in both rear footpads.

### Evaluation of antigen acquisition by flow cytometry

Antigen acquisition was evaluated using fluorescently labeled ovalbumin (ova) as previously described^8–11, 16^ Ovalbumin (A5503, Sigma-Aldrich) was decontaminated of lipopolysaccharide using a Triton X-114 detoxification method and tested with Pierce LAL chromogenic endotoxin quantitation kit (88282, Thermo Fisher Scientific). Ovalbumin was labeled using an Alexafluor 488 succinimidyl ester labeling system (A20100, Thermo Fisher Scientific). CHIKV-infected mice were inoculated with 20 µg AF488-labeled ovalbumin via intramuscular injection into both calf muscles (10 µg per calf) 24 h after virus inoculation, and popliteal LNs were collected 2 days later for analysis of ova^+^ LNSCs by flow cytometry. As positive controls, mock-infected mice were inoculated with 20 µg AF488-labeled ovalbumin and 10 µg polyI:C via intramuscular injection into both calf muscles (10 µg ova-488 and 5 µg polyI:C per calf).

For preparation of single-cell suspensions for flow cytometry, the left and right popliteal LNs were combined for each mouse (2 LNs/sample), minced in Click’s media (Sigma-Aldrich) with 22G needles (Exelint), and digested for 1 hour at 37°C in 94 μg/mL DNase I (Roche) and 250 μg/mL collagenase type I and 250 μg/mL collagenase type IV (Worthington Biochemicals). Cell suspensions were passed through a 100 μm cell strainer (BD Falcon), and total viable cell numbers were enumerated by trypan blue exclusion. All single-cell suspensions were incubated for 15 minutes at 25°C in LIVE/DEAD Fixable Visolet Dead Cell Stain (Invitrogen) to identify viable cells and then stained for 45 minutes at 4°C with anti–mouse FcγRIII/ II (2.4G2; BD Pharmingen) and the following antibodies from BioLegend diluted in FACS buffer (PBS with 2% FBS): anti-CD45 (30-F11), anti-CD31 (clone 390), and anti-PDPN (8.1.1). Cells were washed 3 times in PBS/2% FBS and then fixed for 15 minutes in 1× PBS/1% PFA and analyzed on a BD LSR Fortessa cytometer using FACSDiva software. Analysis was performed using FlowJo software (Tree Star Inc.) and GraphPad Prism version 10.1.1. Data were evaluated for statistically significant differences using a 1-way ANOVA test followed by Tukey’s multiple-comparison test. *P* < 0.05 was considered statistically significant.

### Phosphorothioate oligonucleotides and conjugation to protein

Oligonucleotides were synthesized by Integrated DNA Technologies (IDT) with phosphorothioated oligonucleotides at every linkage as previously described^11^. Oligonucleotides were conjugated to ovalbumin as previously described^11^ using iEDDA-click chemistry^45^. GeoMX tags: DNA tags were provided by IDT using standard desalting and 5’ Amino Modifier C6 with 70 phosphorothioate bonds. GeoMX Barcode 1 target sequence: 5’/5AmMC6/GTTAGGAGGGTCCTTCTAATGTTAACGCCCGAATATTAGTCATATTTTGCTAGCGCCTATCAGCGTAAGA-3’ ; GeoMX Barcode 2 target sequence: 5’/5AmMC6/ GGCGATCCAGCCGGTATACCTTAGTCACATATACTATCGTAATATTGGCGGTTGCTGACAAGTAAATACG-3’. psDNA tags for single cell sequencing: psDNA2 5’/5AmMC6/AGACGTGTGCTCTTCCGATCTNNNCCTGAATTCGAGNNNGCTCACCTATTAGCGGCTAAGG/3Bio/ and psDNA4 5’/5AmMC6/AGACGTGTGCTCTTCCGATCTNNNTCAGGTACCTGANNNGCTCACCTATTAGCGGCTAAGG/3Bio/ Endotoxin levels were quantified using the Pierce Limulus Amebocyte Lysate Chromogenic Endotoxin Quantitiation kit (ThermoScientific) to be less than 0.5EU/mg for either ovalbumin or ovalbumin conjugated to psDNA. Protein concentration was measured using the Pierce BCA Protein Assay kit (thermo scientific) and confirmed on a protein SDS gel with silver stain.

### Preparation of single-cell suspensions

21 or 42 days following vaccination with 1E4 PFU of VV-WR and 20 μg of ova-psDNA per footpad, popliteal lymph nodes were harvested from 13 mice, with 4 lymph nodes mechanically separated with 22-gauge needles in 2 mL of EHAA media. Mouse popliteal lymph nodes were split into two digestion media’s. Eight popliteal LNs were digested in collagenase D (1mg/mL) and DNase 1 (0.25mg/mL) for 30 min at 37°C as described previously^9^ for isolation of the dendritic cells. The other 18 popliteal LNs were digested with 0.25 mg of Liberase DL (Roche, Indianapolis, IN) per mL of EHAA media with DNAse 1(Worthington, Lakewood, NJ) at 37°C to isolate the stromal cell population for 1 hr at 37°C with pipetting every 15 min to physically agitate the digested tissues. The stromal cell digestion media was replaced with fresh digestion media after 30 min. Following digestion, cells were filtered through a screen and washed with 5 mM EDTA in EHAA + 2.5% FBS (R&D Systems, Cat. No.: S12450). The 8 popliteal LNs digested for dendritic cells isolation were then stained with CD11c (N418), CD11b (clone #M1/70) and B220 (clone #RA3-6B2), and a live/dead dye (BD Biosciences Cat. No.:564406). Live cells were then sorted into four tubes on a FACS Aria Cell Sorter (BD): sorted CD11c-APC Cy7 (Clone N418 1:400)+ cells, sorted CD11b PE-Cy7 (Biolegend clone M1/70)+ cells, sorted B220 PE (Clone RA3-6B2)+ cells and Fixable Viability Stain 510 (BD Biosciences Cat. No.:564406) ungated live cells, which were recombined at a 4:4:1:1 ratio, respectively. For the remaining lymph nodes in the stromal cell digest, cells were stained with CD45-PE (clone # 30-F11) and Ter119-PE (clone # TER-119) followed by magnetic bead isolation using the Miltenyi bead isolation kit. CD45-negative cells that passed through the column were then washed. Both sorted and selected (CD45+ and CD45-) cells were then washed with PBS in 0.1% BSA as described in the Cell Prep Guide (10x Genomics) and counted using a hemacytometer. Final concentration of cells was approximately 1600 cells/µL and approximately 10–20 µL were assayed.

### Single-cell library preparation using the 10x Genomics platform

Cells were assayed using the 10x Genomics single-cell 3’ expression kit v3 or v3 HT kit according to the manufacturer’s instructions (CG000053_CellPrepGuide_RevD) and CITE-seq protocol (cite-seq.com/protocol Cite-seq_109213) with the changes outlined in^11^ which include cDNA amplification and cleanup and amplification of antigen tag sequencing libraries. All libraries were sequenced on an Illumina NovaSeq 6000 with 2 x 150 base pair read lengths.

### Preparation of Spatial Transcriptomics Slides

The popliteal lymph nodes were harvested either 3 or 6 weeks after immunization and infection with 20ug ova-psDNA containing a Nanostring GeoMx spatial transcriptomics barcode tag and 1e4 pfu per foot of vaccinia virus. Lymph nodes were fixed for 24-hours in 10% phosphate buffered formalin, and then placed in 70% Ethanol until embedding. The lymph nodes were embedded in paraffin wax, and (7 uM) sections were cut. Slides were stained with Lyve-1-647 (Clone: EPR21771), CD11c-Biotin (Clone: D1V9Y), CD4 (Clone: 4SM95). The nuclei were counterstained with SYTO™ 82 Orange Fluorescent Nucleic Acid Stain (Cat. No.: S11363). Biotinylated antibodies were visualized with Streptavadin-AF594 (Cat. No.: S32356) and the CD4 antibody was visualized with OPAL 520 (Akoya Biosciences, Cat. No.: FP1487001KT). The ROI’s selected to undergo GeoMx® DSP spatial transcriptomics were chosen based on Lyve-1 staining, CD11c staining and CD4 staining. Regions were selected based on the size of the area or the maximum size allowed for the R01.

### scRNA-seq gene expression processing

FASTQ files for the day 2 and day 14 time points were processed as previously described. FASTQ files for the day 21, day 42, and dual immunized samples were processed using the Cell Ranger count pipeline (v6.0.1). Initial filtering of gene expression data was performed using the Seurat R package (v4.4.0). The CD45+ and CD45-samples were processed separately and were combined into separate Seurat objects that included all immunization conditions (2d, 14d, 21d, 42d, dual immunized). Cells were filtered based on the number of genes expressed (>250 and < 5000) and the percentage of mitochondrial reads (<20%). Genes were filtered to only include those detected in >5 cells. Gene expression reads were normalized by the total mouse reads for the cell, multiplied by a scale factor of 10,000, and log-transformed (NormalizeData). Normalized mouse counts were scaled and centered (ScaleData) using the top 2,000 variable features (FindVariableFeatures). The scaled data were used for PCA (RunPCA) and the samples were integrated using the Harmony package. The integrated data were then used to identify clusters (FindNeighbors, FindClusters) and run UMAP (RunUMAP).

For the CD45-samples, we used the integrated cell clusters to generate an initial set of broad cell type annotations using the R package clustifyr (v1.12.0)^21^ and reference data from Immgen^23^. These annotations were checked for accuracy and further refined using known cell type markers including Ptprc, Pdpn, Pecam1, Cd3e (T cells), Cd19 and Cd79a, (B cells), and Cdh1 and Krt8 (epithelial cells). To annotate LEC subsets, endothelial cells were filtered, re-integrated using the Harmony package^46^, and re-clustered. LEC subsets were classified using the clustifyr package along with previously published data^14^.

For the CD45+ samples, we used the integrated cell clusters to generate an initial set of cell type annotations using the R package clustifyr (v1.12.0) and combined reference data from Immgen^23^ along with published data for DC subsets. These annotations were further refined using known cell type markers including Ptprc, Nkg7 (NK cells), Cd3e (T cells), and Cd19 (B cells). For visualization and downstream analysis, DCs were then filtered and re-integrated using the Harmony package^47^.

### Quantification of antigen signals

To compare antigen signals between samples, we divided the antigen counts present in each cell by the total number of antigen UMI counts present in the library (counts per million). To account for different levels of background signal present in each library, we calculated the 75^th^ percentile of normalized counts for B and T cells for each library. We then used this value as an estimation of background signal since B and T cells should not take up antigen. The background signal estimated for each sample was then subtracted from the normalized counts for each cell. For visualization and downstream analysis, these background-corrected values were then log1p-transformed.

To identify antigen-high (Ag-high) and -low (Ag-low) cells, we used the calculated antigen scores for cLECs, fLECs, and collecting LECs from the day 14 time point. We used this time point since it is an intermediate time point in our time course and had distinct populations of Ag-high and Ag-low cells. We focused on cLECs, fLECs, and collecting LECs since they are the subsets with the highest antigen levels. For each subset we used k-means clustering to divide cells into two groups based on the day 14 antigen scores. To ensure consistent classification of Ag-high cells for each time point, we used the antigen score thresholds from the day 14 cells to identify Ag-high cells in the day 21 and day 42 time points. For the dual immunized samples, we identified Ag-high cells separately for the 21 day and 42 day antigen scores.

### Prediction of antigen archiving ability and identification of archiving gene modules

To identify gene expression modules associated with antigen archiving, we trained separate random forest classification models using the Ag-high and -low cells identified for day 14 cLECs, fLECs, and collecting LECs. To identify an initial set of features to use for training each model, we identified genes that were differentially expressed in Ag-high cells for at least one LEC subset from at least one of the time points. Differentially expressed genes were identified using the presto R package (https://github.com/immunogenomics/presto) and were filtered for those with an adjusted p value <0.05. Classification models were trained using the Ranger R package (v0.15.1)^48^. Cells from each LEC subset were split into equal training and testing groups. To identify optimal parameters for model training, a grid search was performed by varying the number of trees (num.trees), the number of variables to possibly split at each node (mtry), and the minimum node size to split at (min.node.size). In addition, the ratio of Ag-high and Ag-low cells used for training was also varied by randomly sampling these groups at different ratios. To identify features most predictive of antigen class for each model, features were filtered based on importance using a p-value cutoff of 0.05 and models were retrained. Ag-high and Ag-low gene modules were identified for each trained model by dividing features into two groups based on whether they were up- or downregulated in Ag-high cells. To identify optimal training parameters, model performance was assessed based on F1 score. Model performance was also assessed independently using cells from a single replicate from published data for LNSCs from naive mice^16^. To do this we used each model to predict Ag-high LECs in naive mice and compared expression of the Ag-high and Ag-low gene modules for the predicted Ag-high cells. For each LEC subset, the models were then filtered to select those with the highest F1 score and that showed differential expression of the Ag-high and -low gene modules for predicted Ag-high LECs from naive mice.

The optimized models for each LEC subset were used to predict Ag-high and -low cells in the 21 day, 42 day, and dual immunized samples. Based on the predicted antigen classes, cells were divided into three groups, 1) has high levels of antigen based on quantified antigen scores (Ag-high), 2) has low levels of antigen but is predicted to be Ag-high (Ag-competent), or 3) has low levels of antigen (Ag-low). For the dual immunized mice, cells were divided into four groups, 1) classified as Ag-high for both antigens (double-high), 2) classified as Ag-high for only one antigen (single-high), 3) classified as Ag-low for both antigens but predicted to be Ag-high (Ag-competent), or 4) classified as Ag-low for both antigens (Ag-low). To compare expression of the Ag-high and Ag-low gene modules derived from each model, module scores were calculated using the Seurat package (AddModuleScore). To identify cell processes that could influence antigen archiving, gene ontology terms (biological process and cellular component) were identified for the Ag-high and Ag-low gene modules for each LEC subset using the clusterProfiler package. Ontology terms were filtered to only include those with >10 genes, <500 genes, and a p value <0.05.

Antigen archiving ability was assessed for mock- and CHIKV-infected samples (3 biological replicates each) using previously published data^16, 17^. These data were processed and cell types were annotated as previously described^16^. Ag-competent cells were predicted using the classification models trained using day 14 cLECs, fLECs, and collecting LECs as described above. Antigen archiving ability was assessed for mock-vs CHIKV-infected mice by comparing the fraction of predicted Ag-competent cells for each biological replicate.

### Processing of GeoMx DSP data

Sequencing libraries generated from the Nanostring GeoMx DSP platform were sequenced on an Illumina NovaSeq6000 at a depth of 200 million reads. FASTQ files were processed using the GeoMx NGS Pipeline v2.0 software. Downstream processing and analysis was performed following best practices described by Nanostring using the GeomxTools (v3.4.0) R package (Griswold M, Ortogero N, Yang Z, Vitancol R, Henderson D (2024) GeomxTools: NanoString GeoMx Tools. R package version 3.6.2). Data were filtered to only include segments with >1000 aligned reads, >75% of reads aligned, >80% of reads stitched, >80% of reads trimmed, and >100 estimated nuclei. Probes were filtered to remove outlier probes based on the Grubb’s test and aggregated for each gene. Genes were filtered to remove those detected in <1% of segments. For gene expression data, counts were normalized based on the 75^th^ percentile of counts present in each segment. Antigen counts were normalized based on background signals estimated using counts for negative control probes. To allow the 21 day and 42 day antigen signals to be compared together on the same plot, signals were scaled separately for each antigen (relative Ag signal, Figure 4D). To do this z-scores were calculated for the 21 day antigen signal for all regions from the 21 day and dual immunized mice, and 42 day antigen signal for all regions from the 42 day and dual immunized mice.

## Acknowledgments

We thank the University of Colorado Genomics Core and the Human Immune Monitoring Shared Resource for their assistance with 10x Genomics captures and GeoMx slides, respectively.

## ETHICS APPROVAL AND CONSENT TO PARTICIPATE

This article complies with the current ethical laws of the United States.

## CONSENT FOR PUBLICATION

All authors provide consent for publication of this primary article.

## AVAILABILITY OF DATA AND MATERIAL

All data and material within this article will be available upon reasonable request to the corresponding author.

## CONFLICT OF INTEREST

We declare no competing interest.

## AUTHORS’ CONTRIBUTIONS

RS performed bioinformatic analysis, conceptualized experiments, and drafted the paper, TAD performed experiments, analyzed data, and drafted the paper, CL performed experiments, conceptualized experiments and revised the paper, TF performed experiments and revised the paper, AU performed experiments and analyzed data, JRH conceptualized experiments and revised the paper, TEM conceptualized experiments and revised the paper and BAJ conceptualized experiments, analyzed data, drafted the paper, revised the paper.

## FUNDING

BAT was funded by NIH R01 AI121209, R01 AI155474 and R21 AI155929, a Department of Medicine ASPIRE Award, the University of Colorado Anschutz Medical Campus GI and Liver Innate Immune Programs and the Waterman Family Foundation for Liver Research.

## CONFLICT OF INTEREST

We declare no competing interest.

## Notes

### Competing Interest Statement

The authors have declared no competing interest.

