## Supplemental figures for "A specific and portable gene expression program underlies antigen archiving by lymphatic endothelial cells"

*Sheridan et. al. Supplemental Figures*

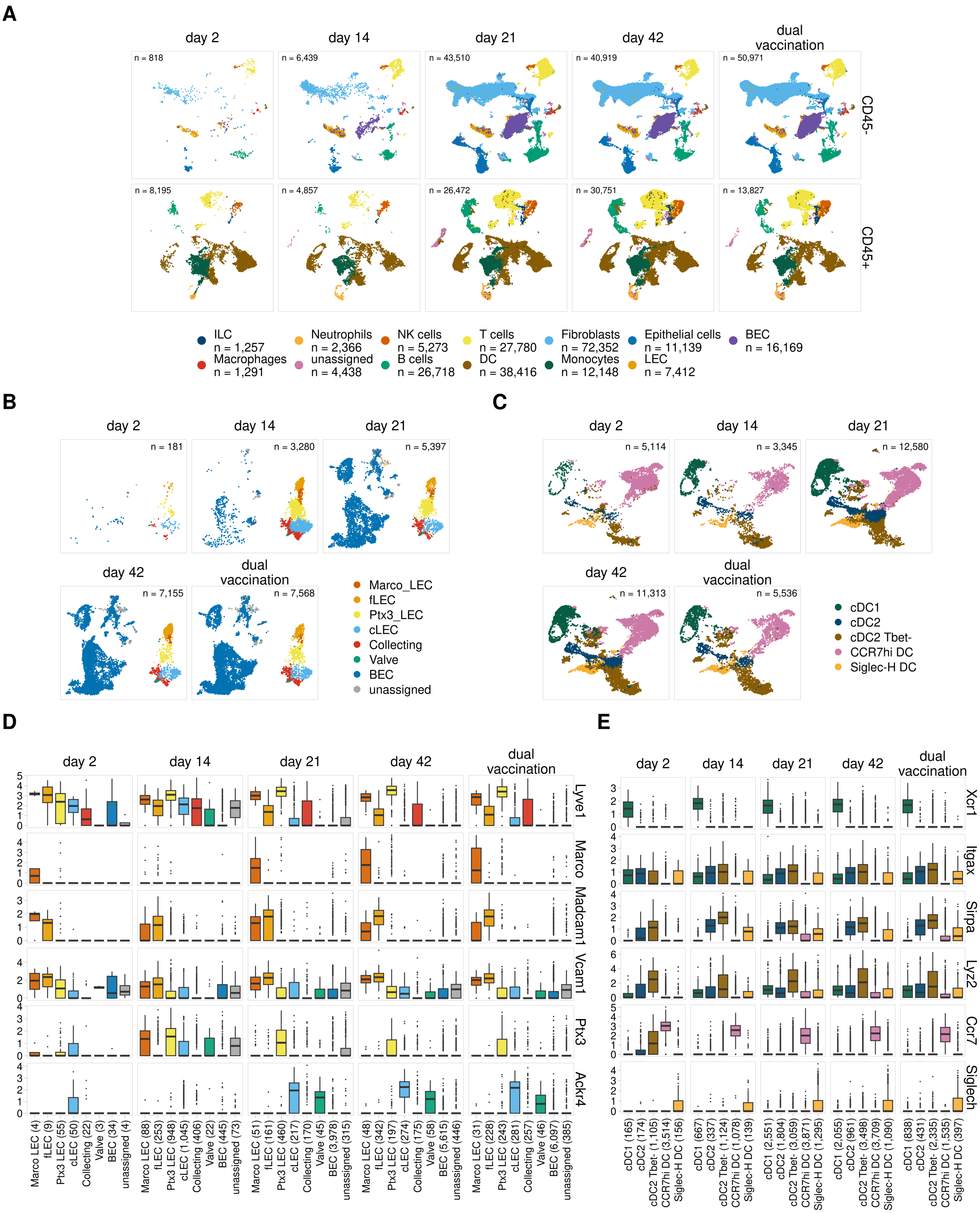


*Figure S1.*

A. *UMAP projections show cell types identified for CD45- and CD45+ samples.*
B. *UMAP projections show LEC subsets identified for each timepoint.*
C. *UMAP projections show DC subsets identified for each timepoint.*
D. *The expression of key marker genes is shown for LEC subsets.*
E. *The expression of key marker genes is shown for DC subsets.*

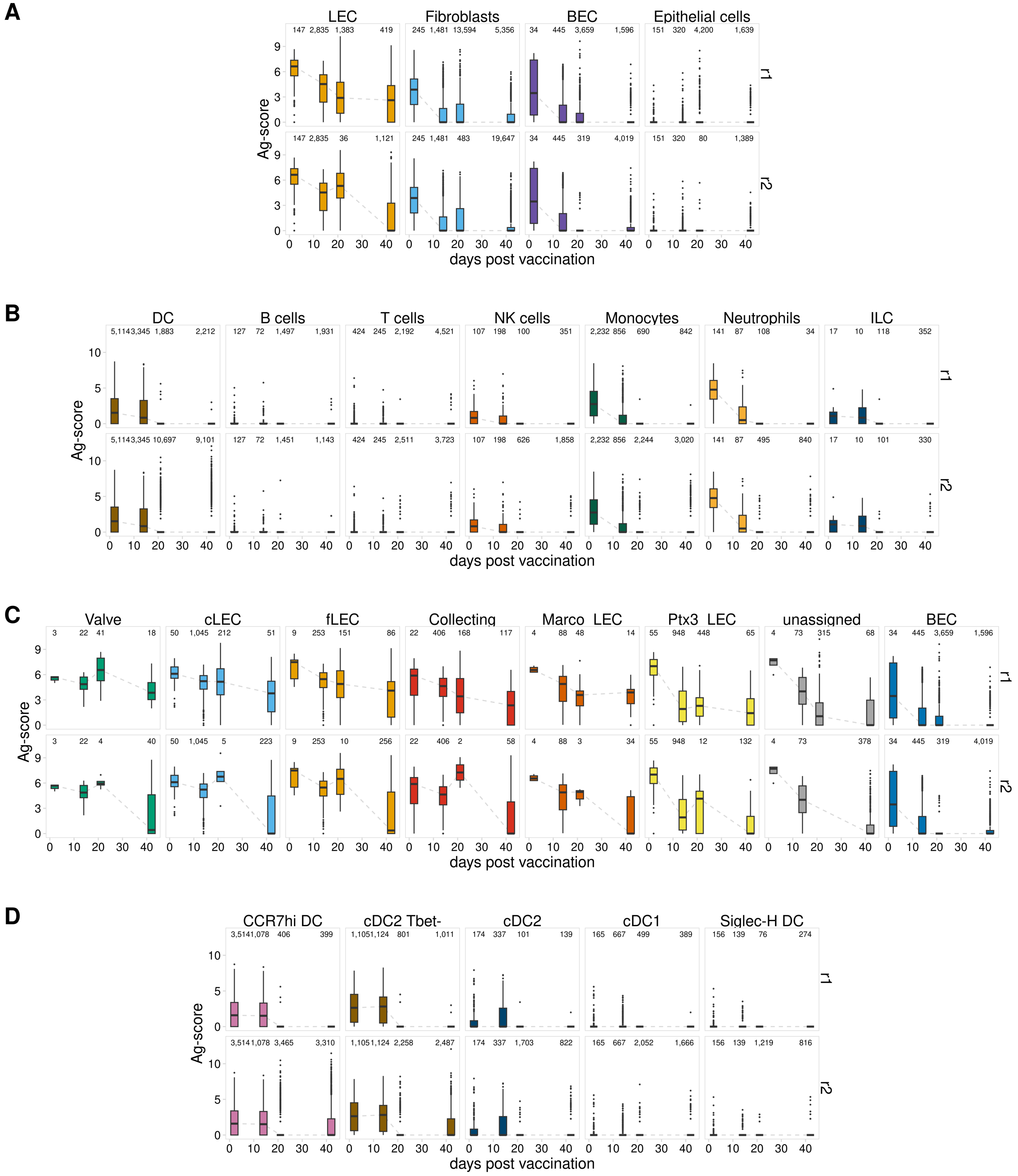


*Figure S2.*

A. *Ag-scores are shown for each timepoint for CD45- cell types. Two biological replicates are shown for the day 21 and day 42 timepoints. The number of cells identified for each cell type is shown above each timepoint.*
B. *Ag-scores are shown for each timepoint for CD45+ cell types, as described in A.*
C. *Ag-scores are shown for each timepoint for LEC subsets, as described in A.*
D. *Ag-scores are shown for each timepoint for DC subsets, as described in A.*

*Sheridan et. al. Supplemental Figures*

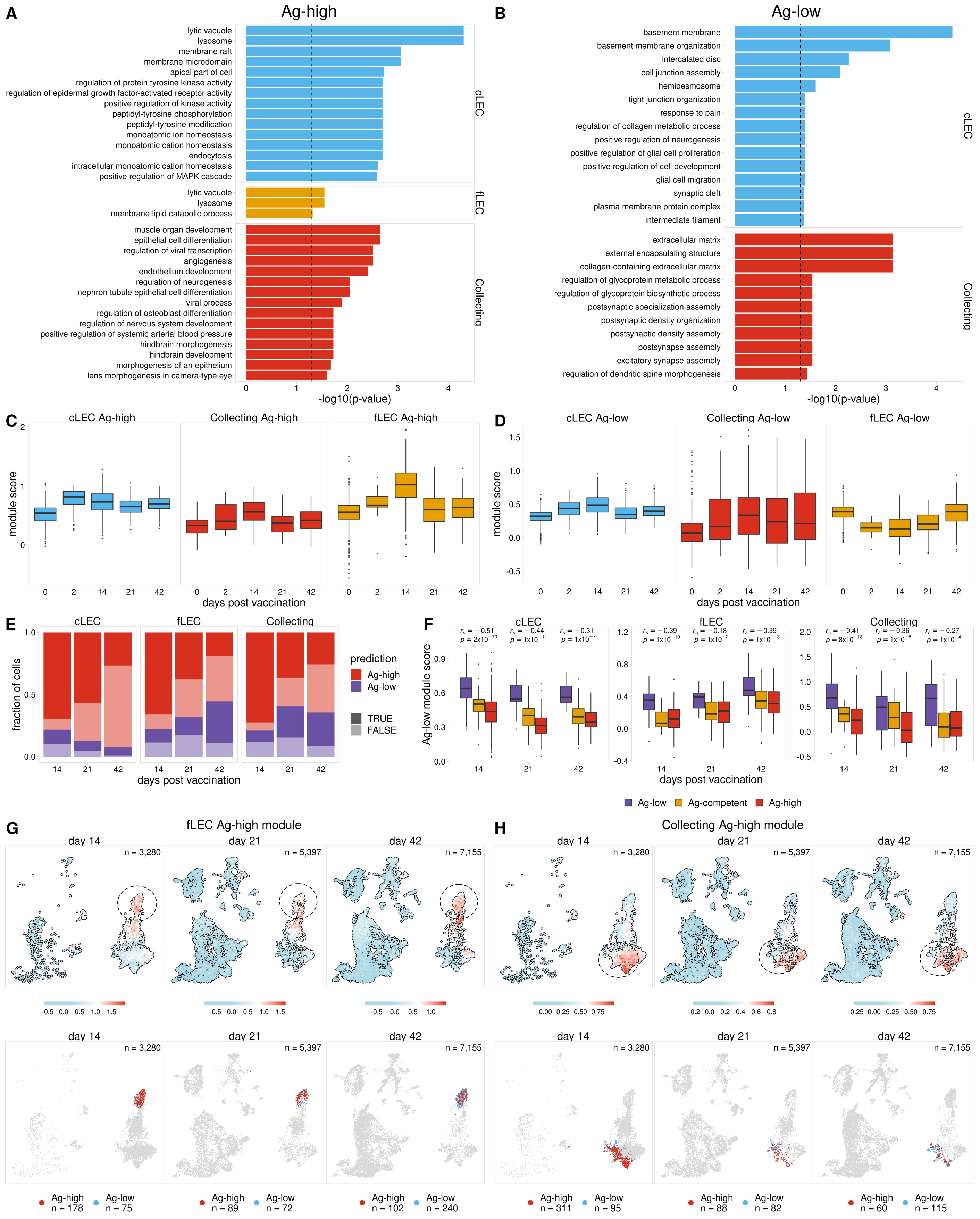

Figure S3.

A. The top 15 gene ontology terms are shown for Ag-high gene modules for each cLECs, fLECs, and collecting LECs.
B. Top gene ontology terms are shown for Ag-low gene modules as described in A.
C. The expression of each Ag-high gene module is shown for naive LECs (0 days post vaccination) along with each time point post immunization.
D. The expression of each Ag-low gene module is shown as described in C.
E. The fraction of cells predicted to be Ag-low and Ag-high is shown for each LEC subset.
F. Ag-low module score is shown for Ag-low, Ag-high, and predicted Ag-competent LECs. The Spearman correlation between Ag classes and Ag-low module score is shown for each timepoint. One-sided p values were calculated and adjusted using Benjamini-Hochberg correction.
G. UMAP projections show fLEC Ag-high module scores for each timepoint.
H. UMAP projections show collecting LEC Ag-high module scores for each timepoint.

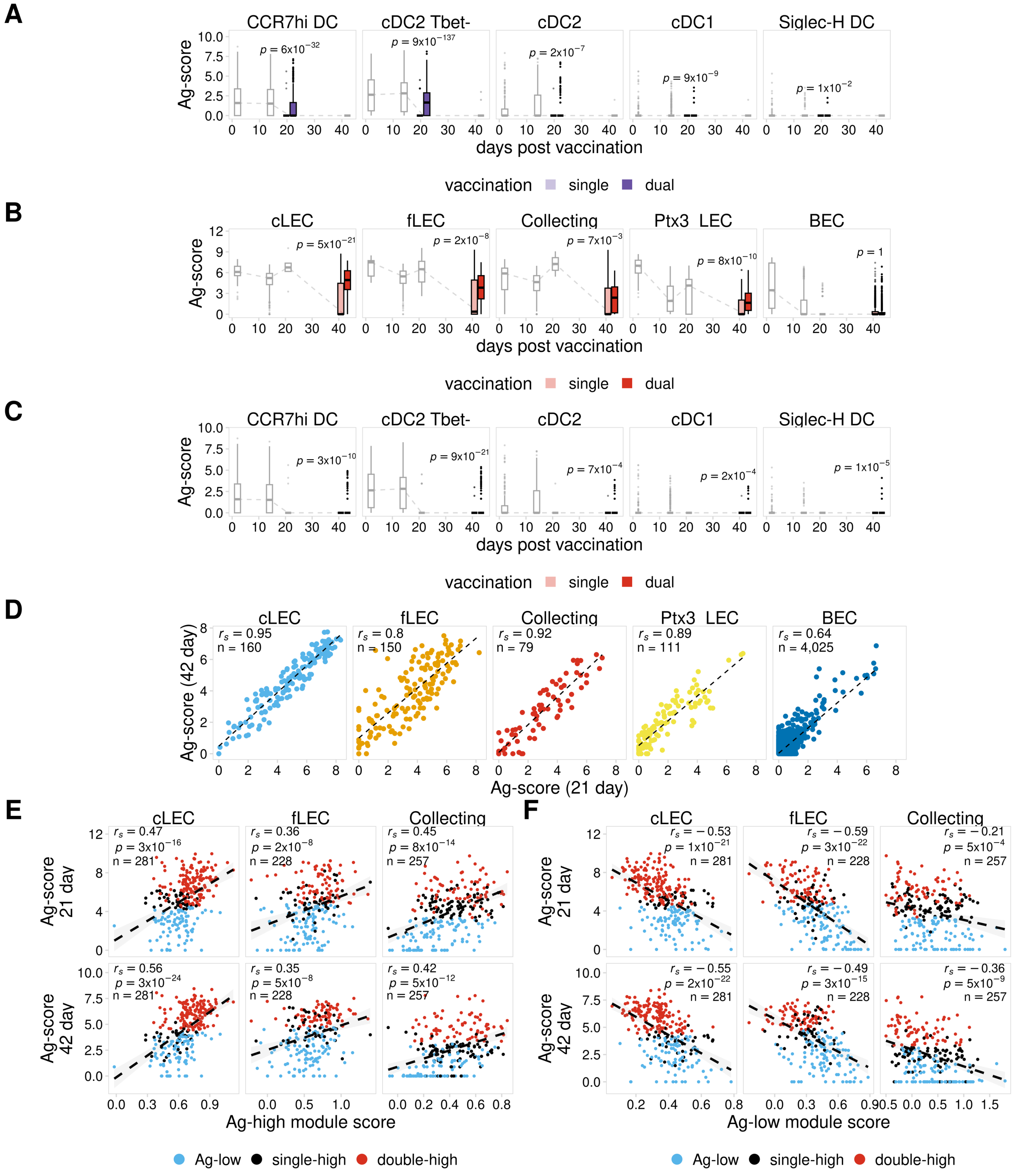


Figure S4.

A. Ag-score is shown for single and dual vaccinations for the 21 day timepoint for DC subsets from a biological replicate. Other timepoints are shown in grey. P values were calculated using a one-sided Wilcoxon rank sum test with Benjamini-Hochberg correction.
B. Ag-score is shown for the 42 day timepoint for LEC subsets from a biological replicate, as described in A.
C. Ag-score is shown for the 42 day timepoint for DC subsets from a biological replicate, as described in A.
D. 21 day and 42 day Ag-score is compared for LEC subsets from a biological replicate.
E. 21 day and 42 day Ag-scores are compared with the Ag-high module scores for each LEC subset. The Spearman correlation is shown for each comparison, p values were calculated and adjusted using Benjamini-Hochberg correction.
F. 21 day and 42 day Ag-scores are compared with the Ag-low module scores as described in E.

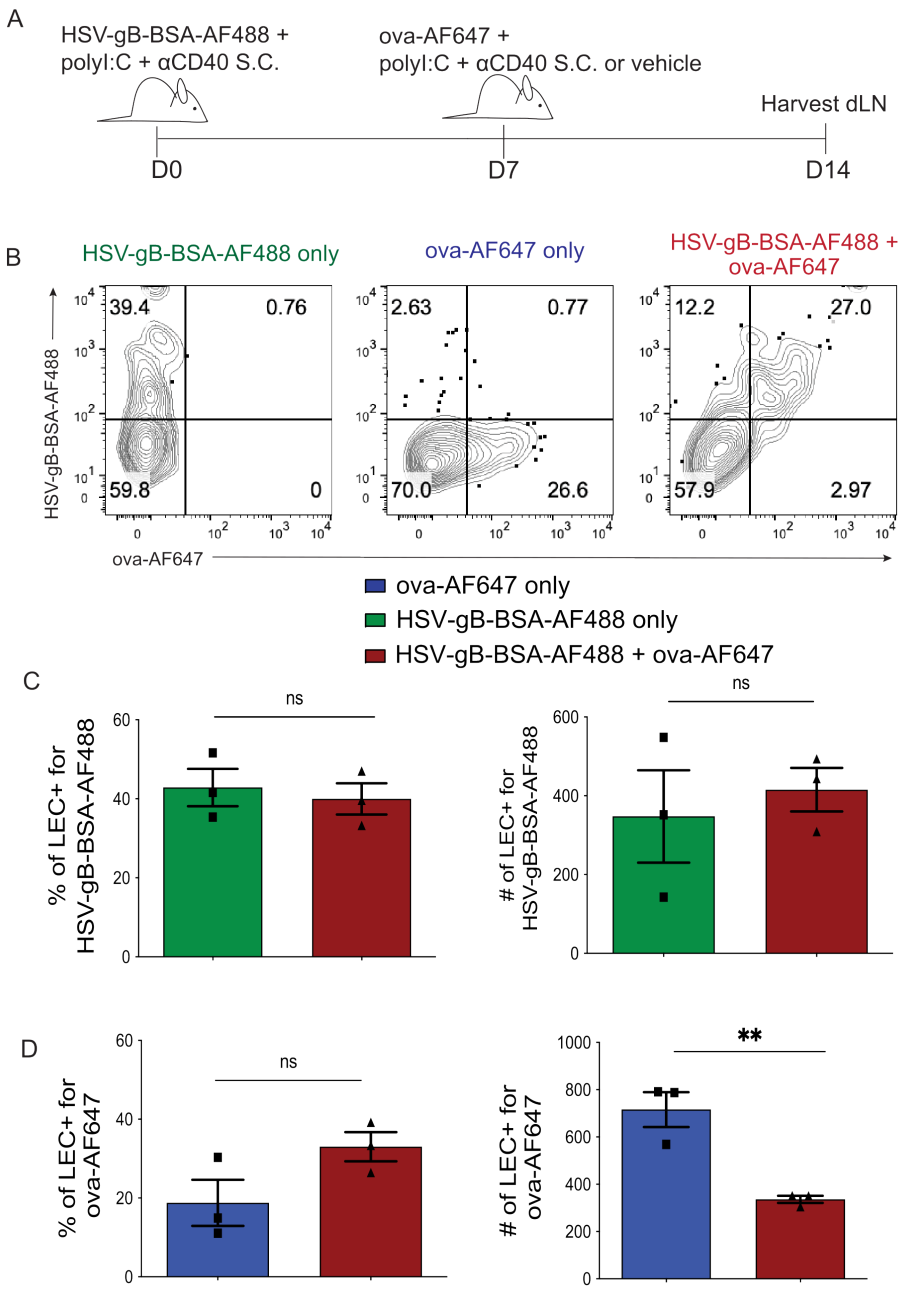


Figure S5. Lymphatic endothelial cells can acquire multiple antigens following sequential immunizations.

A. Mice were immunized subcutaneously with 10 μg of HSV-gB-BSA-AF488, 5 μg of polyI:C, and 5 μg αCD40 in the footpads. After 7 days, mice were immunized with 10 mg of ova-AF647, 5 mg of polyI:C, and aCD40 in the same location. Mice were sacrificed a week later and draining popliteal LN were harvested.
B. Representative flow plots of respective treatments after gating on LEC (CD45-PDPN+CD31+).
C. Quantification of the percent and total number of LEC that are positive for HSV-gB-BSA-AF88 in mice immunized with only HSV-gB-BSA-AF488 (green bar) or mice immunized with HSV-gB-BSA-AF488 and ova-AF647 (red bar).
D. Quantification of the percent and total number of LEC that are positive for ova-AF647 in mice immunized with only ova-AF647 (blue bar) or HSV-gB-BSA-AF488 and ova-AF647 (red bar). Statistical analysis was done using an unpaired t-test where the p-value of the single-immunized mice and dual-immunized mice was <0.0001. Errors bars are mean ± standard error of the mean. In each experiment, at least n=3 mice per group were evaluated and the experiment was repeated n=5 times with similar results. Shown is the data from one of the experiments.

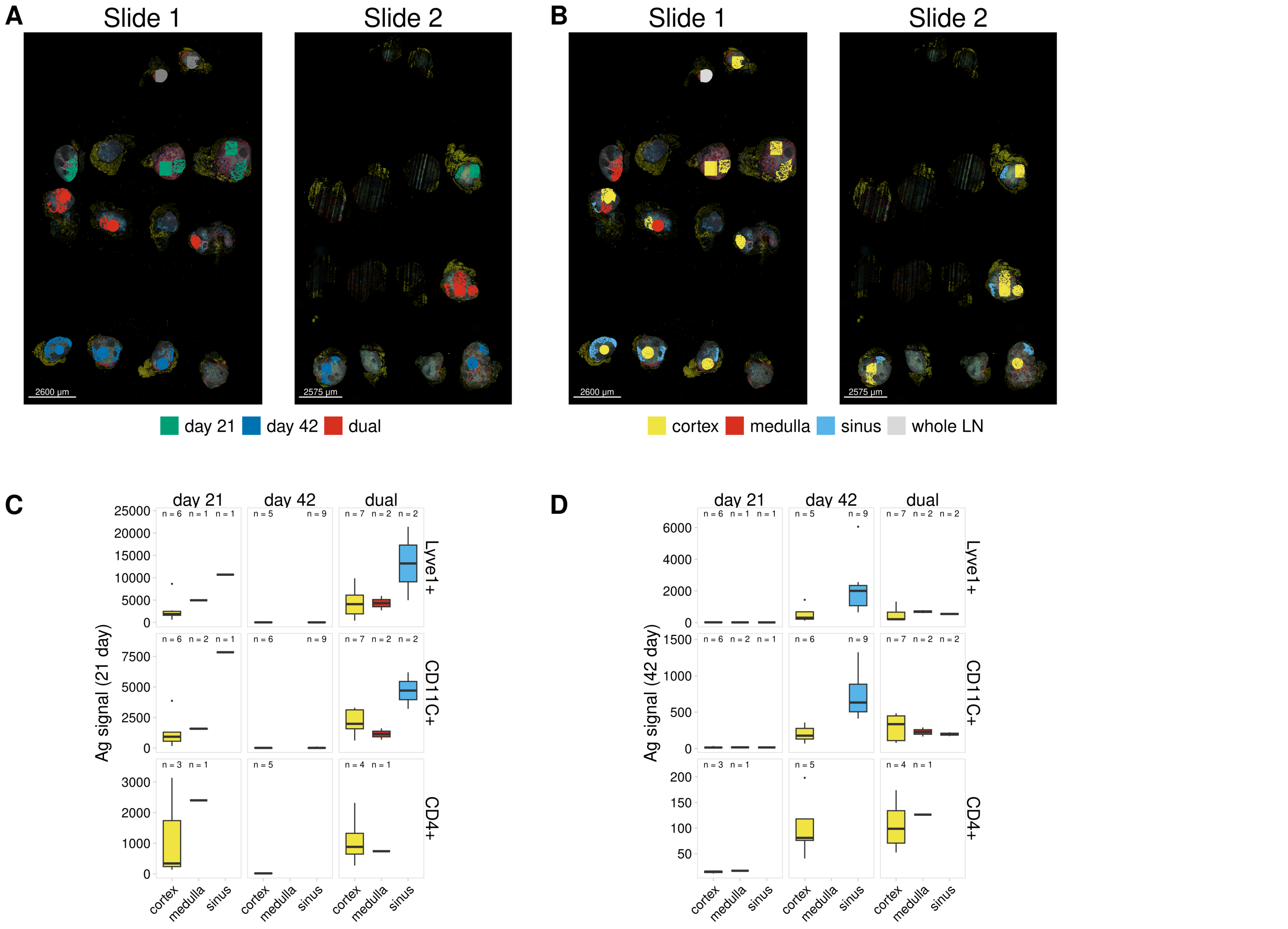


Figure S6.

A. Sample identity is shown for all regions analyzed using the GeoMx DSP platform.
B. Annotated regions are shown as described in A.
C. Normalized 21 day antigen signal is shown for each sample and each region segmented based on Lyve1 (LECs), CD11c (DCs), and CD4 (T cells). The number of plotted segments is shown above each boxplot.
D. Normalized 42 day antigen signal is shown as described in C.

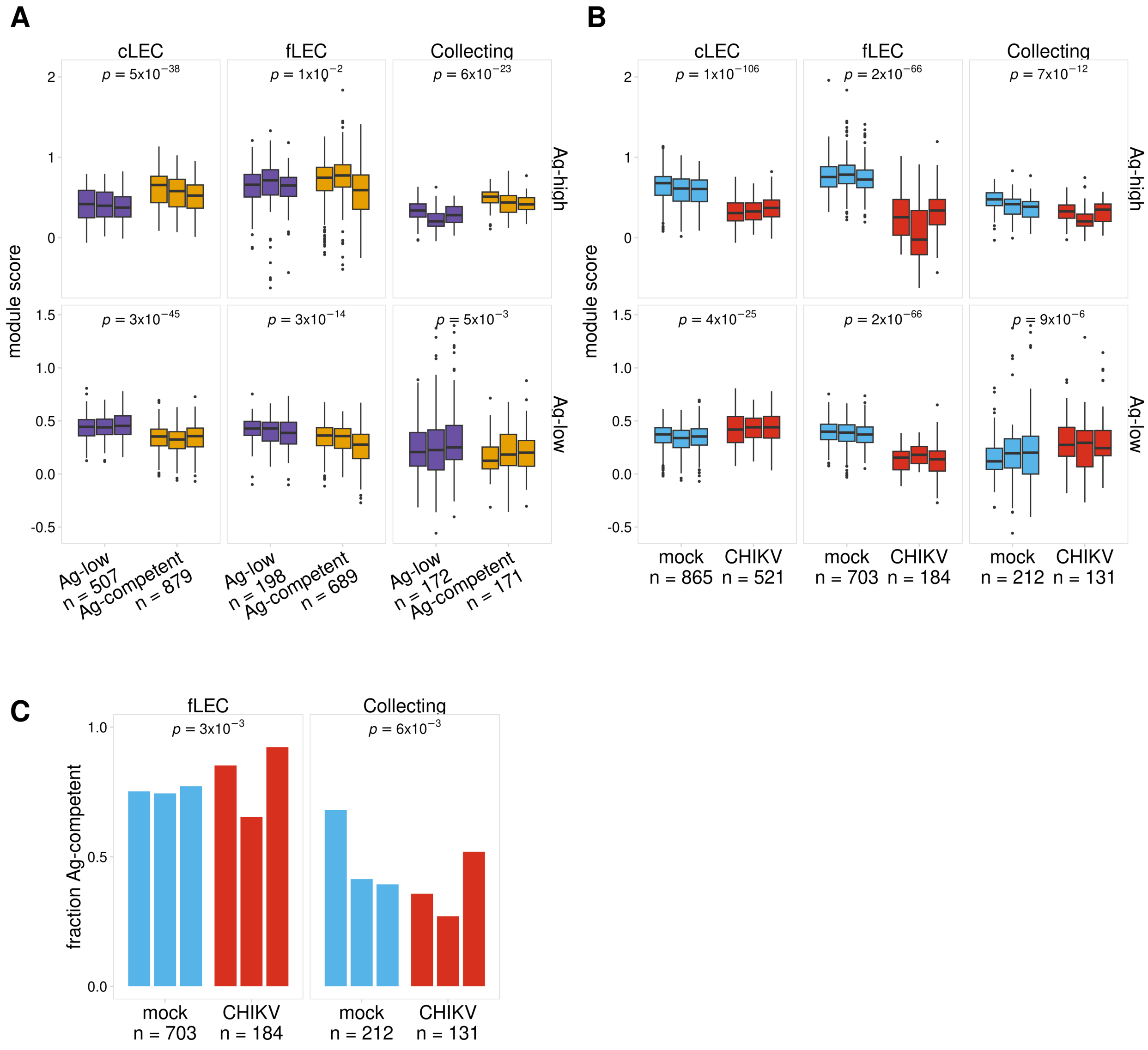


Figure S7

A. Ag-high and Ag-low module scores are shown for the predicted Ag classes. P values were calculated using a two-sided Wilcoxon rank sum test with Benjamini-Hochberg correction.
B. Ag-high and Ag-low module scores are shown for mock and CHIKV-infected mice for each biological replicate. P values were calculated using a two-sided Wilcoxon rank sum test with Benjamini-Hochberg correction.
C. The fraction of cells predicted to be Ag-competent is shown for mock and CHIKV-infected mice for each biological replicate. P values were calculated using Fisher's exact test with Benjamini-Hochberg correction.
